# *TET2* and *TP53* Mutations Cooperatively Modulate the Response to Inflammation to Promote Leukemic Transformation

**DOI:** 10.64898/2026.01.20.700390

**Authors:** Pablo Sánchez Vela, Brianna Kelly, Claudia Brady, Jonathan Foox, Jacob Glass, Richard P Koche, Matthew Wereski, Isabelle Csete, Sebastian El Ghaity-Beckley, Bridget K Marcellino, Ross L Levine, Alan H Shih

## Abstract

*TET2* is a commonly mutated gene in hematologic malignancies, including as an initiating event in clonal hematopoiesis (CH). Its mutation alters hematopoietic self-renewal, differentiation, and systemic inflammation responses. *TP53* mutations co-occur with *TET2* mutations and are also observed in patients with high-risk clonal hematopoiesis and hematologic malignancies. Using a murine model, we found that HSPCs with both mutations initially promoted a myeloproliferative phenotype. Over time these double mutant HSPCs acquire additional genomic alternations, leading to disease progression to acute leukemias including B-ALL. We observed enhanced inflammatory signatures at transformation and identified *NLRP1* as a target of TP53 activation. Decreased response to an inflammatory cell death pathway in the setting of *TP53* mutation allows cells to tolerate inflammatory stress. This pathway also modifies response to chemotherapies that induce protein translational stalling. Our results identify a hematopoietic stem cell stress response pathway with implications on adaptation to inflammation and chemotherapy tolerance.

**Significance:** *TET2* and *TP53* mutations co-operate leading to advanced hematologic malignancy. *TET2* mutations promote an inflammatory environment and *TP53* mutation supports tolerance to this inflammatory stress.

## Introduction

Mutations in *TET2* are amongst the most common across hematologic malignancies including acute myeloid leukemia (AML) (20%) and the precursor condition Clonal Hematopoiesis (CH) (9-10%) (1,2). *TET2* loss of function mutations confer enhanced stem cell fitness and alters the differentiation of myeloid and lymphoid cells (3,4). Mutations most commonly manifest as founder mutations. This is observed in cases of CH where *TET2* mutations can expand and predispose to an increased risk of subsequent leukemia progression (5). *TET2*-mutant CH has also been shown to increase risk of an array of age-associated conditions such as cardiovascular disease due to the promotion of pro-inflammatory states (6).

CH due to *TET2* inactivation alone is insufficient to fully transform hematopoietic stem-progenitor cells (HSPCs) and requires additional cooperating mutational events to progress to overt malignant transformation. We and others have demonstrated that these cooperating events influence the disease phenotype. Cooperating mutations in *FLT3* promote progression to AML, *NRAS* to chronic myelomonocytic leukemia, and *IDH2* to angioimmunoblastic T cell lymphoma (7–9).

Another commonly mutated gene in hematologic malignancies is *TP53*. Similar to *TET2* mutations, *TP53* mutations alone are not sufficient to fully transform HSPCs but often require (and promote) the acquisition or selection of additional mutational events, most notably chromosomal abnormalities. The spectrum of the *TP53*-mutant phenotype is wider than that of *TET2* and frequently involve both myeloid and lymphoid leukemias including B- and T-cell derived malignancies such as large cell lymphomas and acute lymphoblastic leukemias (10,11). *TP53* is found to be mutated in 10% of AMLs, particularly with adverse risk cytogenetics (12). Both WHO classification of hematolymphoid tumors and the International Consensus Classification (ICC) identify *TP53* alteration as distinct subtypes of myelodysplastic syndrome (MDS) and AML (13). These mutations are associated with high-risk cytogenetics and in general poor prognosis due to limited response to the majority of treatment regimens (14). Recent studies also confirm that *TP53* mutations support leukemia development by altering the tumor microenvironment immune response (15). Amongst the mutations that can co-occur with *TP53* but which its transformation effects are less understood is *TET2.* More specifically, the types of leukemia phenotypes it promotes and cell intrinsic inflammatory adaptations required for transformation are yet to be uncovered.

In this study we demonstrate that up to 10% of *TET2* mutant acute leukemia patients carry co-occurring *TP53* mutations. To demonstrate that concurrent *TET2* and *TP53* mutations promote transformation, we developed a genetically engineered mouse model (GEMM) with co-mutation of both of these alleles. This resulted in a myeloproliferative phenotype that evolved to acute myeloid/lymphoid leukemia. Gene expression and cytokine profiling demonstrated activation of inflammatory pathways in HSPC during leukemic transformation. While this occurred, inflammatory cell death, also known as pyroptosis, was impacted by *TP53* loss. *NLRP1*, a component of this pathway, is a target of TP53 transcription and with *TP53* loss, this inflammation-induced pathway is desensitized. NLRP1 acts as a key sensor of cellular stressors, encompassing response to cytotoxic chemotherapy that engages this pathway through ribotoxic translation stress. Manipulation of this pathway may have direct implications in response to standard anthracycline acute leukemia chemotherapy and adaptation in inflammatory stress environments that are promoted by cooperating *TET2* and *TP53* mutations.

## RESULTS

### While *TET2* and *TP53* co-occur in acute leukemia, *TP53* mutational status determines prognosis

In order to understand the mutation pattern of *TET2*-mutant acute leukemias, we analyzed a cohort of patients seen at a single center between 2016 until 2020 who had available next generation sequencing data with a series of panels covering both *TET2* and *TP53* genes (MSK-IMPACT, and MSK-IMPACT-Heme) (16,17) . We identified 115 patients with *TET2* mutant acute leukemias (either AML or ALL) (**Fig. 1A**). The majority of mutations were inactivating either due to frameshift mutation or nonsense mutations as a single nucleotide variant (SNV). The most common *TET2* co-occurring mutations were *DNMT3A* (23%)*, SRSF2* (23%)*, ASXL1* (22%)*, RUNX1* (20%), and *NPM1* (20%). Mutations in *TP53* made up 11% of *TET2-*mutant acute leukemia cases mainly through SNVs and often was associated with additional mutations (**Fig. 1B**). Thus, *TET2* and *TP53* mutations do co-occur and with *TP53* mutation frequency comparable to that in acute leukemias overall. In terms of AML prognosis for the different subgroups, *TET2* mutant AML without *TP53* had better overall survival (median ∼60 mo.), and *TP53* mutant leukemias with or without *TET2* mutation had similar poor survival compared to *TET2* alone (median ∼20 mo.) (**Fig. 1C**). Thus, *TP53* mutations predominate in determining survival response.

**Fig. 1.**
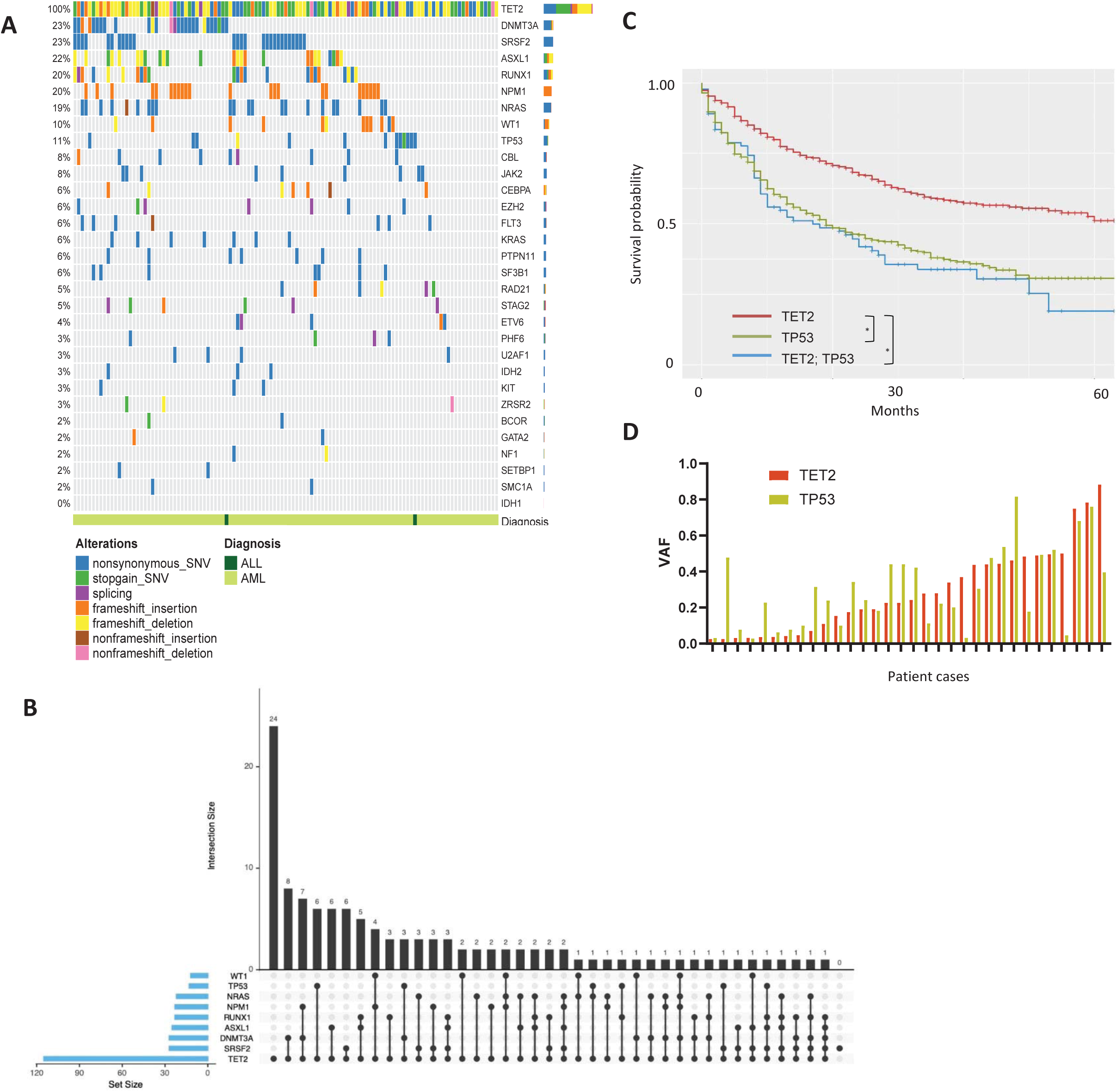
*TET2* and *TP53* mutations co-occur in acute leukemias. **A**, Oncoprint of *TET2* mutant acute leukemias and other co-occurring mutations. *TP53* mutations co-occur at 11% frequency. **B,** Quantification of co-occurring mutations in acute leukemia samples. **C**, Kaplan Meier survival curve of AMLs that are *TET2, TP53*, or double *TET2 TP53* mutant. Log-Rank test for survival of *TET2* mutant AML vs other genotypes. **D**, Hematologic malignancies with co-occurring *TET2* and *TP53* mutations and their associated variant allele frequencies for each patient. * p<.05

Multistep leukemogenesis occurs with sequential somatic mutation acquisition that determine clonal outgrowth trajectories. Clonality is in part implied by variant allele frequency (VAF) -- that originating earlier clonal mutations have higher VAF compared to subsequent gene mutations. We analyzed the VAF composition of our cohort along with select myeloid malignancy patients with these mutations. Of the 33 cases of co-mutation with VAF data, 13 had higher *TP53* mutant VAF, 10 higher *TET2*, and 10 where the VAFs were within a 5% difference (**Fig. 1D**). This suggests that either *TET2* or *TP53* mutation can act as a founder mutation clone or develop earlier, with subsequent secondary mutations. As both mutations can be involved in CH, these findings support the possibility of evolution from either of these founding states.

### Loss of *Tp53* and *Tet2* leads to acute leukemia

To study the potential interactions of *Tp53* and *Tet2* in hematopoiesis we generated a compound mutant mouse by crossing *Tp53^-/-^* germline knockout (KO) mice with *Mx1-Cre^+^ Tet2^fl/fl^* mice, generating *Mx1-Cre^+^Tet2^fl/fl^Tp53^-/-^* (18). In this GEMM upon induction with polyinosinic-polycytidylic acid (PIPC), Cre is expressed that causes floxed exon deletion in *Tet2* resulting in double loss-of-function (*Tp53^-/-^ Tet2^-/-^)* in the bone marrow compartment (**SFig. 1A**). To better study the genetic effects of each genotype in disease evolution, we treated donors from wild-type, single-mutant, and combined mutant mice with PIPC and transplanted them into recipients (n=10). We observed more aggressive disease with combined mutant mice compared to single mutant mice with median survival of ∼125 days (not reached for the other genotypes) (**Fig. 2A**). We further refined and confirmed the disease progression of *Tet2^-/-^Tp53^-/-^* mice with additional transplants (**Fig. 2A**). We assessed the complete blood count of mice at 10 weeks and found that *Tet2^-/-^Tp53^-/-^* mice (TP) compared to WT, *Tet2^-/-^* (T), and *Tp53^-/-^* (P) mice developed anemia (Hct WT 47±2.5%, T 44±3.6%, P 45±3.8%, TP 37±6.3%) and thrombocytopenia (WT 997±232k/µl, T 1111±240k/µl, P 958±196k/µl, TP 402±365k/µl) compared to wild-type and single mutant mice (**Fig. 2B**). White blood cell count was selectively elevated in individual mice (WT 14.4±5.6k/µl, T 15.7±3.4k/µl, P 17.4±4.7k/µl, TP 23.8±23.4k/µl) (**Fig. 2B**).

**Fig. 2.**
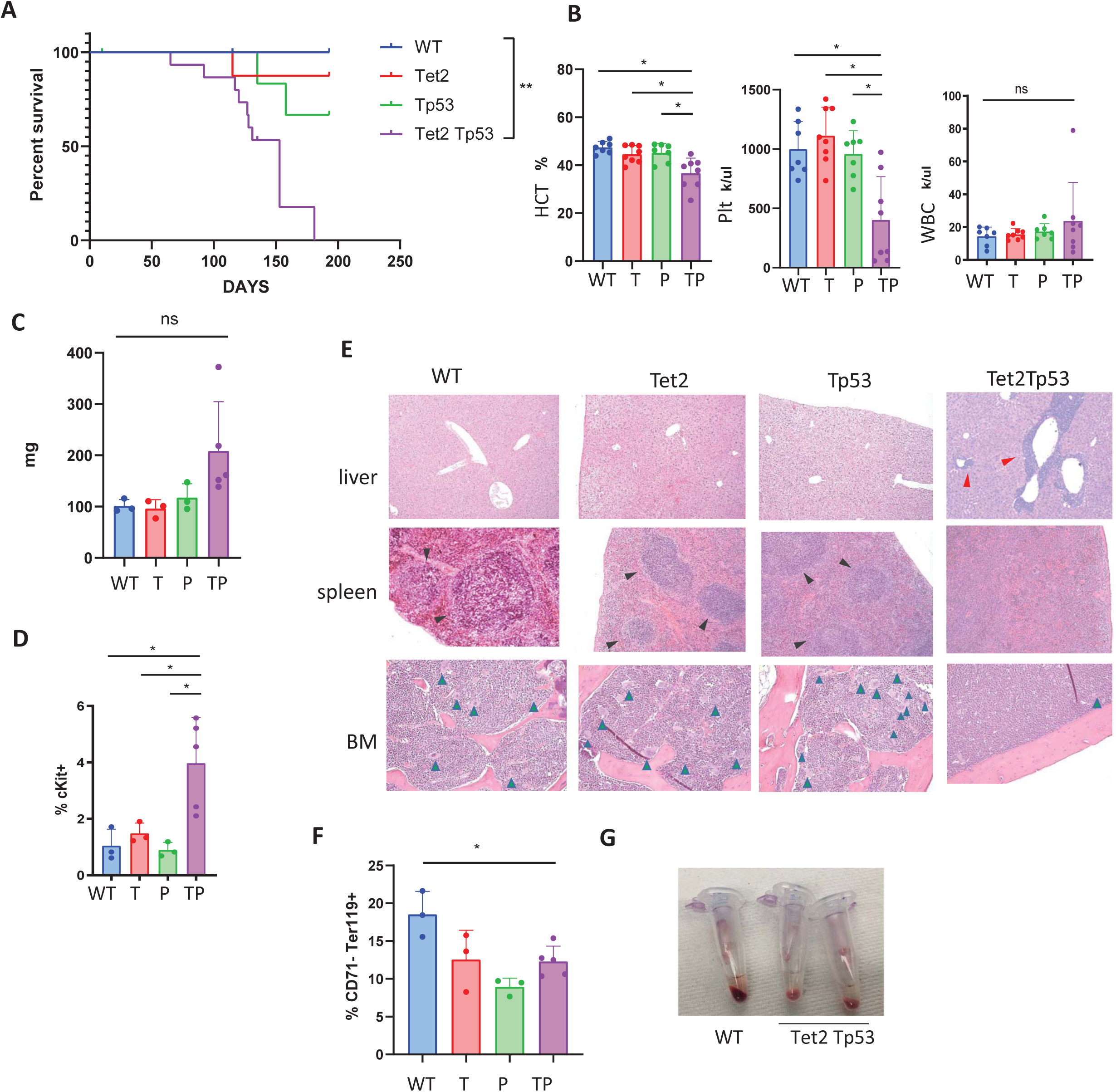
Primary GEMM of *Tet2 Tp53* co-mutations develop hematologic malignancies. **A**, Kaplan-Meier survival curve of mice transplanted with wild type (WT), *Tet2* (T), *Tp53* (P) (n=10) and *Tet2 Tp53* (TP) (n=18) mutant bone marrow (3 mice from each group was censored at 18wks for pathology evaluation). ** p<.01 by Log-rank test for multiple curve comparison. **B**, Peripheral blood count parameters for each transplant group, hematocrit (HCT), platelet count (Plt), and white blood cell count (WBC) at 10 weeks. **C**, Spleen weights at 18wks and time of disease. **D**, Immature cKit^+^ cell percentage in the spleen. **E**, H&E histology sections from liver, spleen and bone marrow (BM) from wild type, *Tet2*, *Tp53*, and *Tet2 Tp53* mutant BMT mice. Red arrow, infiltrating leukemic cells in liver sinusoids. Black arrow, germinal centers in spleen. Green arrow, megakaryocytes in BM. **F**, Bone marrow mature erythroid fraction CD71^-^Ter119^+^ cells. **G**, Gross pathology of bone marrow aspirates with reduced red cell volume in diseased *Tet2 Tp53* double mutant bone marrow. * p< .05, ns- not significant.

Over time *Tet2^-/-^Tp53^-/-^* mice developed more aggressive disease that led to more frank leukemia development. We performed a timed analysis of 3 mice from each group at 18 weeks along with earlier moribund *Tet2^-/-^Tp53^-/-^* mice. *Tet2^-/-^Tp53^-/-^* genotype resulted in splenomegaly (WT 101±12mg, T 96±17mg, P 118±27mg, TP 209±26mg) and increased immature cKit^+^ cells in the spleen (WT 1.0±0.6%, T 1.5±0.4%, P 0.9±0.3%, TP 4.0±1.6%) indicating proliferative disease and secondary hematopoiesis (**Fig. 2C-D**). There was infiltration of hematopoietic cells into the liver, and in the spleen normal splenic architecture was disrupted, germinal centers were no longer clearly formed (**Fig. 2E top & middle**). The bone marrow pathology showed a hypercellular marrow of monotonous cells with reduced megakaryocytes. This reduction in megakaryocytes correlated with the relative thrombocytopenia (**Fig. 2E bottom**). Flow cytometry demonstrated reduced mature CD71^-^Ter119^+^ red blood cells compared to wild-type; similar reduction in mature erythropoiesis was also observed in single mutant mice (WT 18.5±3.0%, T 12.5±3.9%, P 8.9±1.1%, TP 12.3±2.0%) (**Fig. 2F**). In bone marrow aspirates from diseased mice, erythropoiesis was visibly decreased and appeared grossly composed of less red blood cell volume compared to wild type with a paler appearance (**Fig 2G**) in line with the flow cytometric erythroid characterization (**Fig. 2F**). Thus, these mutations account for changes in HSPC differentiation with particular deficits in erythroid and megakaryocytic development.

### Combined *Tp53* and *Tet2* loss leads to both myeloid and lymphoid leukemic disease

In the transplanted mice there was early evidence at 10-12wks of myeloid predominance in the peripheral blood with an increased proportion of CD11b^+^ (Mac1^+^) cells (WT 18.3±1.7%, T 20.9±2.9%, P 20.5±2.4%, TP 39.3±20.5%) compared to lymphoid B-cells (B220^+^); this bias was more pronounced compared to single mutant mice (**Fig. 3A**). Although there was this myeloid bias, we noted variation in individual recipients. We also analyzed the peripheral blood of primary (non-transplant derived) mice at 10 to 12wk time points and observed both lymphoid and myeloid biased differentiation as well (**Fig. 3B**). Specifically, we observed an increased in select mice of the monocytic population CD11b^+^Gr1^-^ that is associated with myeloid bias from *Tet2* mutations (18). In separate select mice, we also observed the expansion of an immature B-cell population that was B220^dim^ (**SFig. 1B**). We have previously reported on *Tet2* mutations impairing B-cell development in germinal centers, and *TP53* deletion or mutation is associated with lymphoid disorders (4,19). Disrupted lymphoid development is also in-line with the known consequences of mutations in these genes. Thus, *Tet2^-/-^Tp53^-/-^* mice initially possess a myeloproliferative phenotype with progressive cytopenias but can also exhibit early signs of lymphoid disease.

**Fig. 3.**
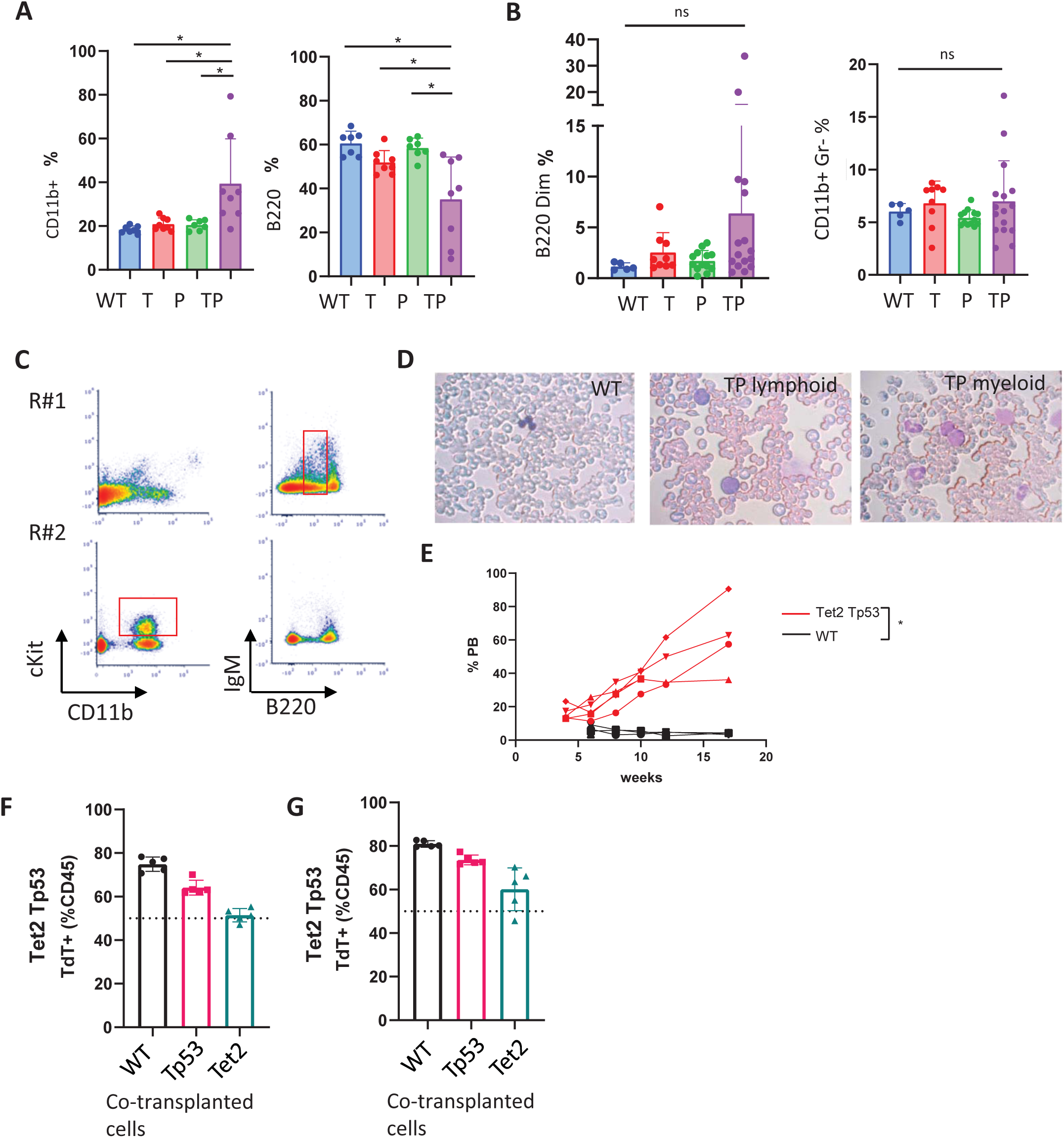
*Tet2 Tp53* mutations in HSPCs cooperate to promote both myeloid and lymphoid phenotypes. **A**, Peripheral blood myeloid CD11b^+^ and lymphoid B220^+^ cell fractions from primary (non-transplanted) mice at 10-12 weeks and **B**, Immature B220^dim^ cells and monocytic CD11b^+^Gr1^-^ cell fractions. **C**, Peripheral blood flow cytometry from two transplanted recipients using the same donor bone marrow displaying independently myeloid CD11b^+^ and lymphoid B220^+^ disease. **D**, Peripheral blood smear from wild type (WT) and lymphoid (L) and myeloid (M) disease. **E**, Peripheral blood WBC fraction of CD45.2^+^ *Tet2 Tp53* and wild type transplanted into recipient mice tracked over time. **F&G**, *Tet2^-/-^ Tp53^-/-^ TdTomato^+^* co-transplanted (and competed) with wild type, *Tp53^-/-^,* or *Tet2^-/-^* bone marrow at (F) 4wk sand (G) 12wks (n=4-5 per group). *p<.05, ns – not significant

In additional transplant experiments (using independent donor mice) we observed both myeloid and lymphoid biased hematopoiesis and leukemias evolved in recipients from the same donor-derived marrow (**Fig. 3C** and **SFig. 1C**). Myeloid disease is associated with CD11b^+^ markers and immature cKit^+^ expression (**Fig. 3C**). In lymphoid disease, B220^dim^ expression was associated with subset IgM^dim^ marker. Peripheral blood smears (**Fig. 3D**) demonstrated blast-like morphology from both cases with enlarged cells and high nuclear – cytoplasmic ratios. In myeloid disease, immature myeloid forms can be observed in the smear with hypo-segmented myeloid precursors. Once more advanced disease was established, myeloid or lymphoid biased disease remained stable upon serial transplantation (**SFig. 1D**). Subsequent recipient mice retained myeloid or lymphoid phenotype. Thus, mutations of *Tet2* and *Tp53* together are likely insufficient to fully transform HSPCs and additional genomic or epigenomic events might be required for transformation. Nonetheless, together they contribute to both myeloid and lymphoid leukemic disease potential.

### Combined *Tp53* and *Tet2* loss leads to stem cell growth advantage

We examined the effect of combined effect of *Tp53* and *Tet2* loss on hematopoietic cell expansion in vivo. Bone marrow *Tet2^-/-^Tp53^-/-^* cells were collected from mice prior to acute leukemic disease. We transplanted these cells, which were derived in a CD45.2^+^ background or wild-type bone marrow cells in a CD45.1^+^ background. By tracking this surrogate marker, we observed competitive growth advantage with expanding CD45.2^+^ *Tet2^-/-^Tp53^-/-^* population over time while wild-type transplanted cells remained stable (WT 4.2±1.0%, *TP* 36±5% at 10wks) (**Fig. 3E**). Thus, the combined mutations provided a stem cell growth advantage compared to WT.

We also asked whether the combined mutant cells had growth advantage over single mutant cells. To this end, we crossed a Td Tomato (*TdT*) fluorescent marker allele into *Mx1-Cre^+^Tet2^fl/fl^Tp53^-/-^* mice (20). We then performed similar transplantation experiments as above using pre-leukemic cells, mixing a 50:50 ratio of *Tet2^-/-^Tp53^-/-^ TdT*^+^ cells with wild-type, *Tp53^-/-^* or *Tet2^-/-^* bone marrow. We then tracked the *TdT^+^* fraction over time. Double mutant cells again out-competed wild-type bone marrow. They also out-competed *Tp53^-/-^* single mutant cells at 4wks and 12wks (**Fig. 3F**). At 4wks *Tet2^-/-^* cells were of equal composition but at 12wks, double mutant cells held an advantage (12wks TdT^+^ vs WT 81±1.5%, vs T 60±9.8%, vs P 73±2.2%) (**Fig. 3G**). Thus, each mutation contributes to the growth advantage in the bone marrow and combined mutations having greater competitive advantage, providing positive selection for these mutations if they co-occur.

### Tet2 Tp53 disease progression is associated with differential methylation and genomic instability

Since TET2 acts as a de-methylase, we hypothesized that methylation changes may be responsible for changes in the disease phenotype. We isolated HSPCs (lineage^-^ Sca1^+^ ckit^+^, LSK) from our initial transplant lymphoid leukemic *Tet2^-/-^Tp53^-/-^* mice and performed methylation analysis. Using reduced representation bisulfite sequencing, we mapped changes in methylation and compared to wild-type LSK cells. These changes were then mapped to differentially methylated regions (DMRs), associated genes, and gene elements. DMRs were able to segregate mutant cell types based on methylation differences (**Fig. 4A**). These methylation differences occurred throughout the gene structure including at CpG islands, promoters, and gene bodies (**Fig. 4B**) but only tended slightly towards hypermethylation as would be expected with loss of *TET2* de-methylase activity (**Fig. 4C**). However, the genes associated with methylation changes did not correlate well with changes in gene-expression and did not readily explain a *Tet2*-methylation based disease mechanism – that hypermethylation of direct target genes was responsible for transcriptional repression (**Fig. 4D**). We mapped the gene expression differences with associated gene methylation differences assed in this study and other previously studied models, *Flt3^ITD^, Tet2^-/-^,* and *Tet2^-/-^ Flt3^ITD^* and the *Tet2^-/-^ Tp53^-/-^* methylation pattern did not significantly better correlate with the gene expression changes (**Fig. 4D**) (7).

**Fig. 4.**
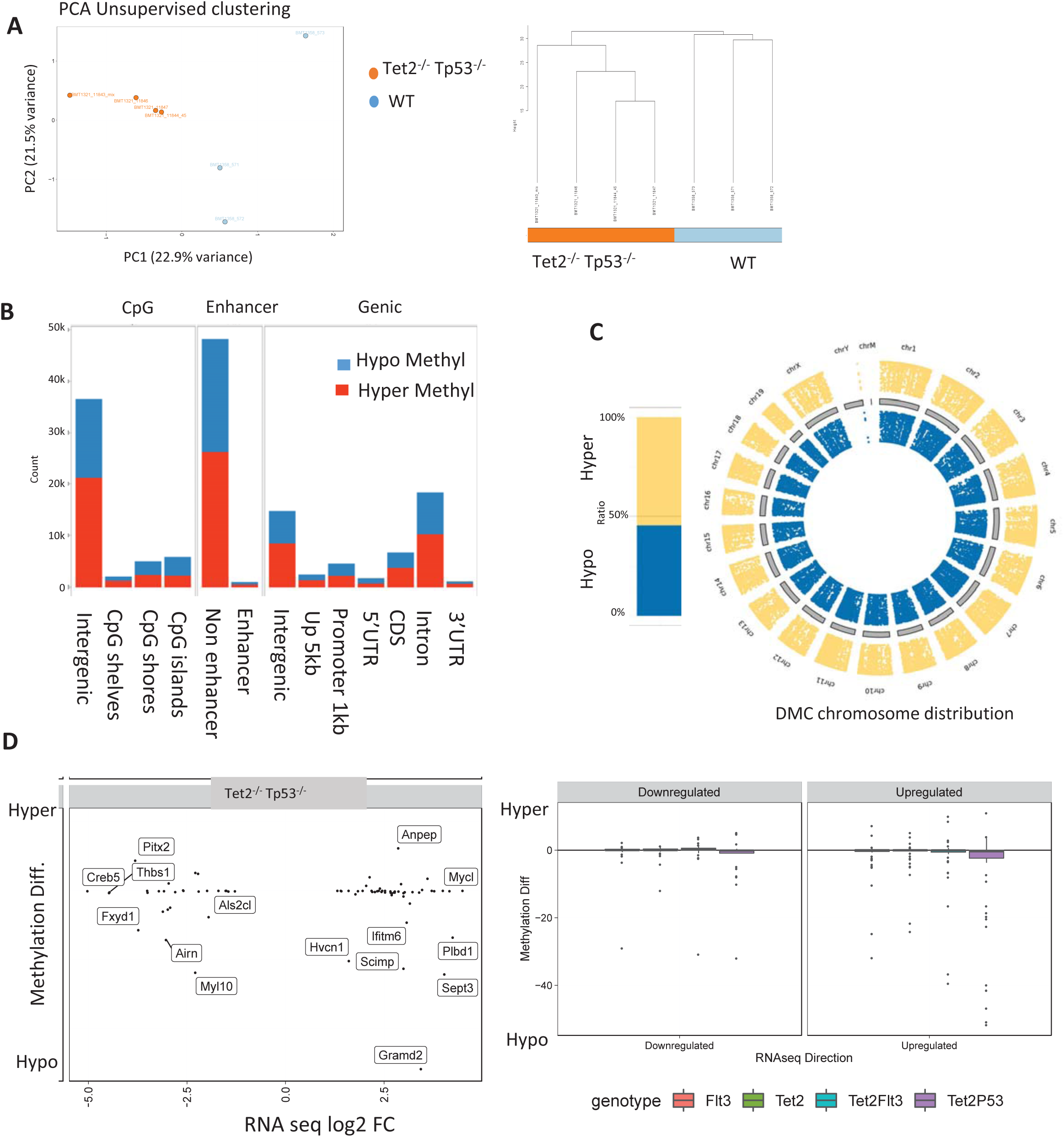
Methylation reduced representation bisulfite sequencing (RRBS) analysis from *Tet2 Tp53* mutant diseased cells. **A**, (left) Principal component analysis (PCA) from RRBS data of methylation in stem-progenitor lineage^-^Sca^+^ckit^+^ (LSK) cells, wild-type and *Tet2 Tp53* mutant. (right) Hierarchical clustering of RRBS profiles from wild type and *Tet2 Tp53* mutant cells. **B**, Hypo- vs Hyper-methylation of specific gene segments assigned to CpG islands (CpG), Enhancer regions, and Gene regions- intergenic, upstream 5kb, promoter to 1kb, 5’UTR, coding sequence (CDS), introns, and 3’UTR. **C**, Analysis of Hypo- vs Hyper-methylation in genomic DNA of *Tet2 Tp53* double mutant vs wild-type and as determined across chromosomes. CpGs filtered to padj < 0.01 & methylation difference >= 25% or <= -25%, Hypermethylated differentially methylated cytosines (DMCs): 26,759, Hypomethylated DMCs: 22,573 by methylKit. **D**, Association between differentially expressed genes and (left) methylation level in *Tet2 Tp53* mutant leukemia and (right) divided into downregulated and upregulated and as associated with respective gene’s methylation level in *Tet2, Flt3, Tet2 Flt3,* and *Tet2 Tp53* mutant models relative to WT.

We hypothesized that instead additional genetic alterations may determine disease evolution. We performed murine targeted capture-based sequencing from diseased mice (lymphoid and myeloid) to discern specific genetic changes that may cause disease evolution and phenotype. Although select SNVs were observed, we did not observe recurrent mutations at a sufficient tumor level VAF that would readily explain disease evolution (**STable 1**). However, because of the depth of coverage in exome sequencing, we were able to identify copy number variations (CNVs) based on relative depth of reads (**Fig. 5A** and **SFig. 2**). Both myeloid and lymphoid leukemias had gains and losses of chromosome segments compared to *Tet2^-/-^Tp53^-/-^* mice prior to transformation. Thus, genomic instability may be a means of disease evolution. Specifically, patterns of CNV were different between lymphoid versus myeloid disease and these genetic changes along with possible SNVs may drive disease progression and phenotype (e.g. CNV on chromosome 14 were more common in lymphoblastic leukemia). Instead of methylation changes related to *TET2* loss of function, *TP53* loss appears to drive the evolution of leukemia by enabling chromosome instability.

**Fig. 5.**
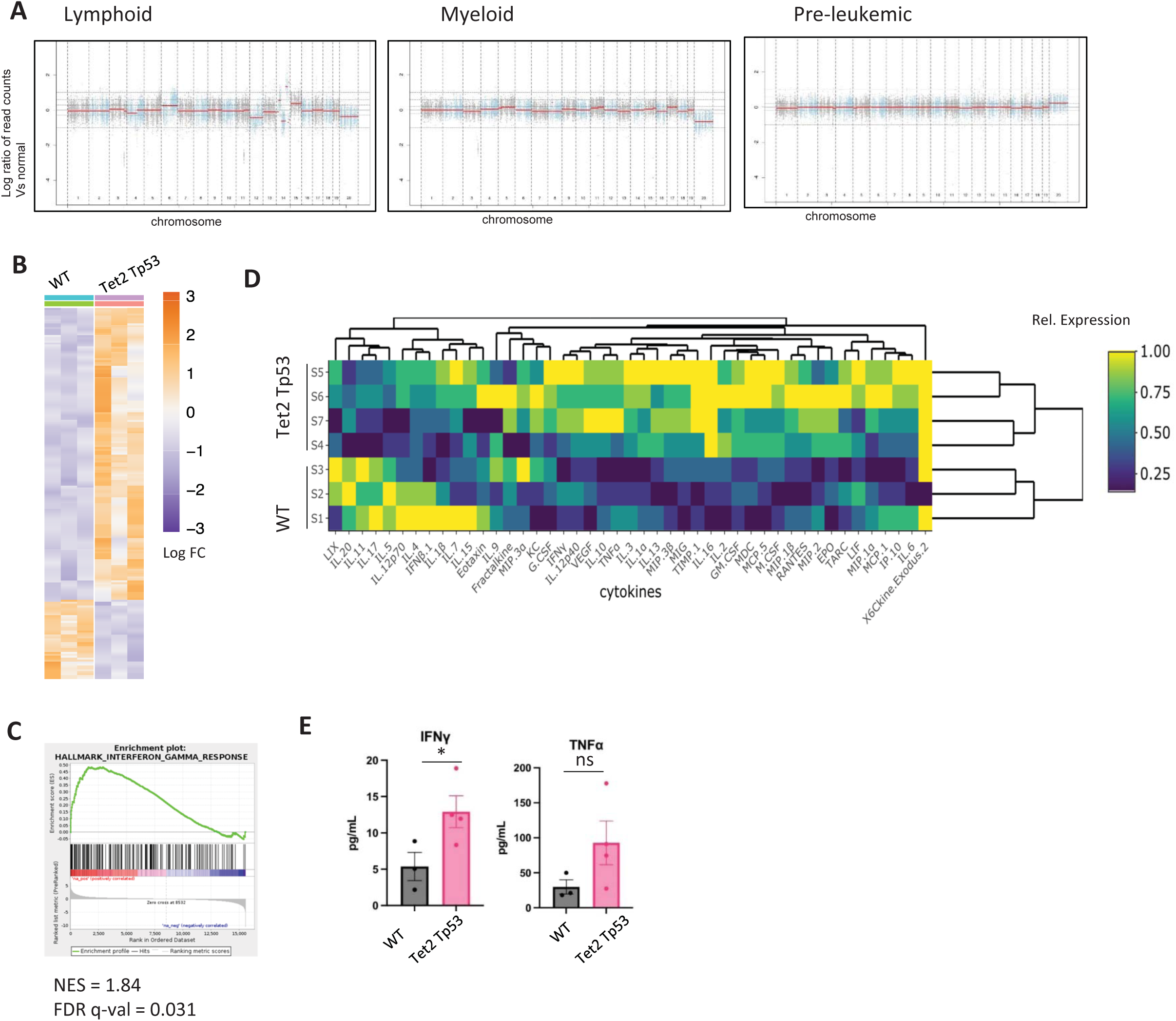
Genomic and proteomic profiling of *Tet2 Tp53* double-mutant disease. **A**, Manhattan plot of copy number variability based targeted exome sequencing coverage of genomic DNA in lymphoid and myeloid diseased mice and in pre-leukemic *Tet2 Tp53* mutant mice. Red line indicates the computed average of the copy number fold over normal in a given segment. **B**, Heat map of differentially expressed genes (RNA-seq) of sorted lineage^-^ Sca^+^ cKit^+^ (LSK) stem and progenitor bone marrow cells. (n=3 biological replicates per group, p-adj <0.5, log2 fold change >1). See also Supplemental Table 2. **C**, GSEA analysis of the differential gene expression signature, interferon gamma response (FDR 0.031, NES 1.85). **D**, Heat map of Luminex-based plasma profiling from wild type (n=3) and *Tet2 Tp53* double-mutant leukemic diseased mice (n=4), and **E**, plasma levels of IFNγ and TNFα. * p<.05, ns- not significant.

### Combined *Tp53* and *Tet2* loss leads to interferon gamma transcriptional signaling in HSPCs upon lymphoid leukemia progression

To further investigate the mechanism of disease progression we performed RNA sequencing on purified bone marrow lineage^-^ Sca^+^ ckit^+^ (LSK) stem progenitor cells at lymphoid acute leukemia stage and compared to wild-type LSKs. *Tet2^-/-^Tp53^-/-^*mutant LSKs had a differential gene expression pattern that distinguished them from wild-type HSPCs (**Fig. 5B** and **SFig. 3A, STable 2**). Using GSEA analyzing associated gene signatures, immune-related signatures were amongst the highest ranking (**SFig. 3B**). Of particular interest was an interferon gamma response signature that was upregulated in the mutant cells (**Fig. 5C**). By qPCR, we confirmed the expression of *Ifng* and interferon-target genes were elevated in bone marrow cells from independent *Tet2^-/-^Tp53^-/-^*mutant leukemic mice compared to wild-type (**SFig. 3C**).

To investigate this inflammatory signature, we profiled the plasma of transplanted *Tet2^-/-^Tp53^-/-^* leukemic (n=4) and wild-type mice (n=3). We found cytokine profiles did segregate the two groups independently (**Fig. 5D** and **STable 3**). *Tet2^-/-^Tp53^-/-^* had several significantly elevated cytokines compared to normal controls. Notably, INFγ and TNFα were elevated, cytokines commonly implicated in inflammatory response and immune modulation (mean, IFNγ WT 5.4 pg/mL, TP 12.9 pg/mL; TNFα WT 29.7 pg/mL, TP 92.7 pg/mL) (**Fig. 5E**). The combination of these cytokines is known to influence inflammatory response and stimulate a pyroptosis cell death pathway and activation of the inflammasome (21). We also noticed upon extraction of the bone marrow that select *Tet2^-/-^Tp53^-/-^* marrows were less cellular compared with wild-type or single mutants (**Fig. 6A**). This was associated with reticulin staining in the bone marrow of disease mice, demonstrating the development of fibrosis that is associated with elevated inflammation but not in other non-inflammatory models (**SFig. 3D**) (22).

**Fig. 6.**
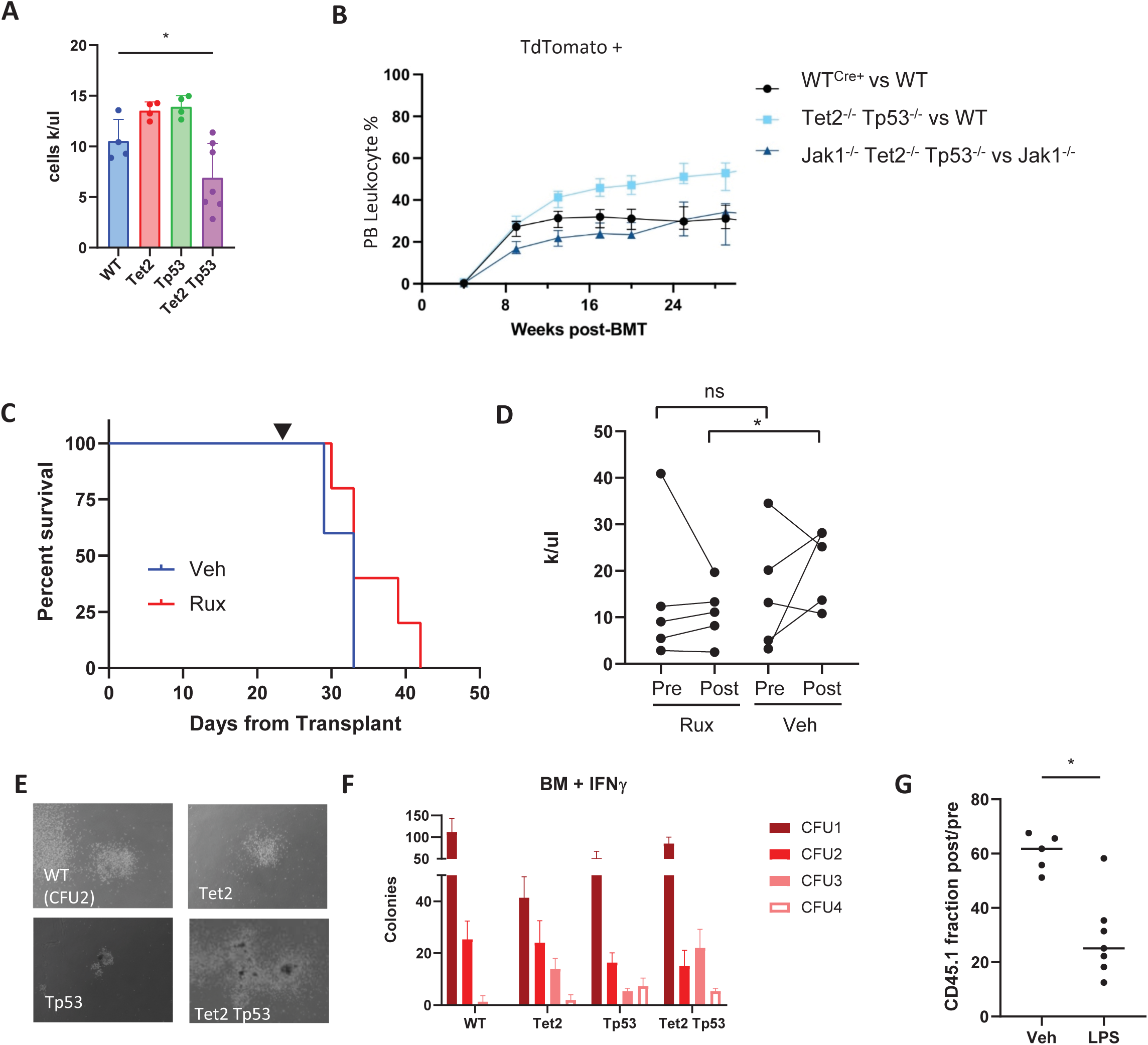
*Tet2 Tp53* co-mutated malignancies develop a pro-inflammatory and inflammation-resistant phenotype. **A**, Relative cellularity from femurs from individual wildtype or mutant mice eluted into equal volumes. **B**, Fraction of TdTomato^+^ (TdT^+^) cells in the peripheral blood. Black circles, WT (TdT^+^) vs WT. Blue square, *Tet2 Tp53* double mutant (TdT^+^) vs WT. Green triangle, *Tet2 Tp53 Jak1* triple mutant (TdT^+^) vs *Jak1* mutant. **C,** Kaplan-Meier survival curves of *Tet2 Tp53* double-mutant diseased mice (BMT) treated with either vehicle or ruxolitinib (60mg/kg daily) (n=5 per arm). (p=0.15) **D**, WBC counts pre- and post-ruxolitinib. **E&F**, Colony forming assay using cytokine supplemented methocult (Stem Cell Technology M3434) treated with IFNγ 50ng/mL. (E) representative colony images and (F) colony number following serial re-platings. **G**, *In vivo* advantage of *Tet2 Tp53* double-mutant disease cells in the setting of inflammation. *Tet2 Tp53* double mutant cells were mixed with wild type CD45.1^+^. The retained CD45.1^+^ fraction was measured between subsequent bleeds after LPS treatment twice a week (2.5mg/kg) for 3 weeks (n=5 Veh, 7 LPS). * p≤.05, ns- not significant.

Since inflammatory signaling occurs through JAK/STAT signaling, we asked whether this differential response to cytokines was important for its growth advantage. We crossed our *Mx1-Cre^+^Tet2^fl/fl^Tp53^-/-^Tdt^+^* mice with a conditional *Jak1^fl/fl^* allele (23). JAK1 phosphorylation from receptor complexes is important for signal transduction and response to cytokine stimulation. For example, by deleting *Jak1*, signaling by INFγ is significantly limited (24). We asked whether *Tet2 Tp53* mutant cells would still retain growth advantage compared to wild-type cells if both cell types were deficient in JAK1-mediated signaling. In competitive transplant experiments, combined mutant cells that had deleted *Jak1* compared similarly to *Tet2^WT^ Tp53^WT^ Jak1^-/-^* cells (**Fig. 6B**). That is, when these cells were transplanted together in recipient mice, *Tet2^-/-^ Tp53^-/-^Jak1^-/-^* cells did not out compete *Jak1^-/-^* cells. In the setting of cytokine signaling defect, the *Tet2 Tp53* combined mutations no longer expanded. Thus, this JAK1 signaling pathway is important for *Tet2^-/-^ Tp53^-/-^* mutation growth advantage.

To test this in a pharmacologic fashion, we transplanted *Tet2^-/-^ Tp53^-/-^* leukemic cells into recipient mice and treated with vehicle or ruxolitinib, a JAK1/2 inhibitor (25). Mice treated with ruxolitinib had a trend towards improved survival, but they still succumbed to disease (**Fig. 6C**). Ruxolitinib was more likely to stabilize or decreased the white count compared to vehicle, suggesting that cytokine signaling is one part of the disease transformation mechanism (**Fig. 6D**). This inflammatory state then may play a role in disease advancement and provides a point of therapeutic intervention.

### Combined *Tp53* and *Tet2* loss leads stem cell advantage in setting of INFγ growth suppression

INFγ signaling is known to suppress hematopoietic stem cell function (26). Exposure reduces stem cell renewal and differentiation, and in patients can contribute to cytopenias during an infection or following CAR T-cell therapy (27). Although we observed a hyper-inflammatory signature with leukemic transformation, we hypothesize that tolerance to these cytokines precedes this at the pre-leukemic HSPC stage to allow for increased cytokines to evolve. To test the effect of INFγ on stem cell activity, we performed colony formation assays and serial re-plating using in vitro semi-solid media, using non-leukemic mice. Wild-type colonies extinguished after 2 re-plating cycles in the setting of INFγ but *Tet2^-/-^*, *Tp53^-/-^*, and *Tet2^-/-^ Tp53^-/-^* bone marrow cells can re-plate for further cycles, although with fewer colonies compared to vehicle treatment (**Fig. 6E-F, SFig. 4A**). This is also true in the setting of combined INFγ and TNFα exposure (**SFig. 4B**). Although single mutant *Tet2^-/-^* and *Tp53^-/-^* mutant cells were able to re-generate colonies more than wild-type, *Tet2^-/-^ Tp53^-/-^*cells most robustly and consistently re-plated in the setting of cytokine suppression (**SFig. 4B**). INFγ continues to be suppressive for mutant HSPCs but they demonstrate in vitro renewal advantage compare to wild-type.

To test for this advantage *in vivo*, we transplanted *Tet2^-/-^ Tp53^-/-^* and wild-type cells into recipient mice. Once cells were engrafted, we treated mice with vehicle or LPS, an agent that induces inflammatory response, including INFγ and TNFα cytokines (**SFig. 4C**). While *Tet2^-/-^ Tp53^-/-^* cells as expected had a competitive growth advantage, with LPS treatment this advantage was even greater (**Fig. 6G**). We tracked the percentage of wild-type CD45.1^+^ and leukemic fraction CD45.2^+^ over consecutive bleeds. In the LPS-treated setting after 3 weeks, this loss of wild-type fraction and increase in leukemia occurred at a faster rate compared to vehicle controls (wild-type fraction retained between bleeds, Veh 60±6.8%, LPS 29±15%).

### TP53 regulates expression of *NLRP1*

When we examined the cytokine profiles of *Tet2^-/-^ Tp53^-/-^* leukemic mice compared to controls, we noted that IL-1β was surprisingly not significantly elevated (mean, WT 17.5 pg/mL, *Tet2^-/-^ Tp53^-/-^* 16.6 pg/mL) (**Fig. 7A**). IL-1β is a cytokine that is produced during inflammatory stress by caspase 1 activation and correlates with inflammatory cell death, pyroptosis. We also noted that although INFγ signaling gene signature was detected in the RNA-seq differentially expressed genes, NOD-like receptor family, pyrin domain containing 1a (*Nlrp1a*) was lower in expression in leukemic mice compared to controls. In *Tp53* WT cells, inflammatory cytokine can induce expression of *Nlrp1a* in both wild type and *Tet2^-/-^* bone marrow (**SFig. 5A**). *NLRP1* is part of the signaling pathway that is activated by pathogens, stress, and cellular damage and its activation leads to the production of IL-1β (28). We confirmed the reduced *Nlrp1a* expression by qPCR from *Tet2^-/-^ Tp53^-/-^* and *Tp53^-/-^*mutant bone marrow cells (**SFig. 5B**). Notably *Nlrp1a* is linked to *Tp53* as the two genes are proximate to each other on chromosome. This synteny of the two genes is preserved in human *NLPR1* and *TP53*, as they also are closely linked (chr17p13.2 *NLRP1* and chr17p13.1 *TP53*). Deletion of *TP53* that occurs in acute leukemias can also confer a copy loss of *NLRP1*. We tested this by performing genomic qPCR quantification on *TP53* deleted AMLs comparing levels of *NLRP1* and *TP53* to a control sequence, and indeed deletion of *TP53* is often accompanied by *NLRP1* deletion (**SFig. 5C**). Although there is no evidence for haploinsufficiency for *NLRP1*, there are cases of homozygous deletion of *TP53* and thus this would be another means of losing this sensor.

**Fig. 7.**
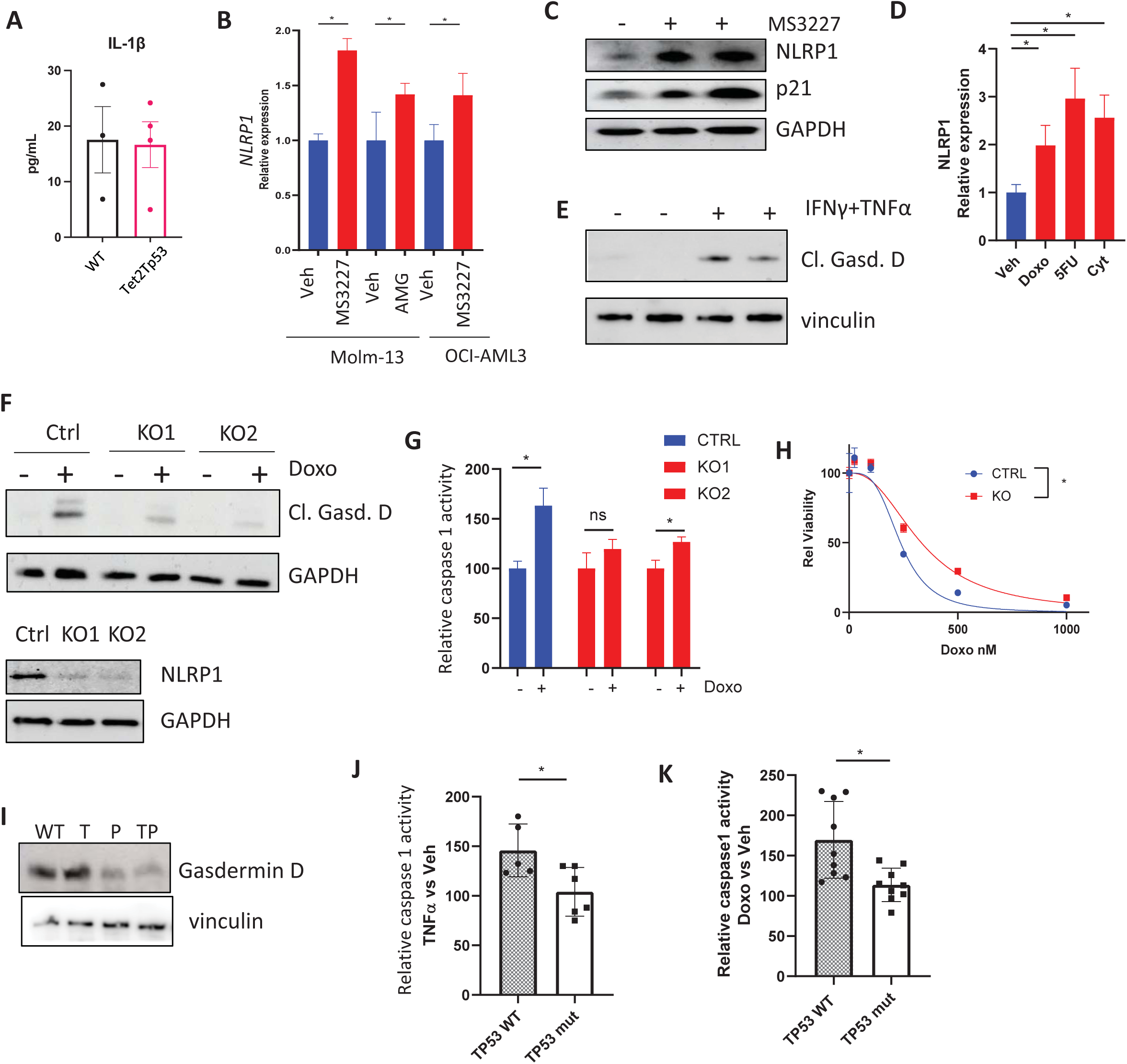
TP53 activates NLRP1 and *TP53* mutations disrupt activation of inflammatory pathway attenuating pyroptosis. **A**, Plasma IL-1β levels in *Tet2 Tp53* mutant disease and wild type mice. **B**, qPCR of the relative *NLRP1* mRNA levels in cell lines treated with MDM2 inhibitor for 24hr (AMG232 or MS3227 250nM). **C**, Western blot of NLRP1, p21, and GAPDH protein expression in OCI AML3 cells treated with vehicle or MDM2 inhibitor MS3227 (250nM) 24hr. **D**, qPCR relative expression of *NLRP1* in MV4-11 cells treated with vehicle, doxorubicin (1µM), 5FU (25µM), or cytarabine (25µM). **E**, Western blot of cleaved gasdermin D and vinculin in wild-type mouse bone marrow stimulated with IFNγ + TNFα (50ng/mL). **F**, (Top) western blot of OCI AML-3 cells Control (Ctrl) or knock-out for NLRP1 (KO1, KO2) for cleaved gasdermin D and GAPDH following treatment with doxorubicin (1µM) & (Bottom) western blot for NLRP1. **G**, Relative caspase 1 activity in OCI AML3 control or *NLRP1* KO cells treated with Doxorubicin (1µM). **H**, Dose response curve to Doxorubicin in OCI AML3 control (IC50 236nM) and *NLRP1* KO (332nM) cells. **I**, Western blot for Gasdermin D, cleaved lower band, and vinculin from bone marrow from wild-type and myeloproliferative stage *Tet2^-/-^, Tp53^-/-^,* and *Tet2^-/-^ Tp53^-/-^* mice. **J&K**, Relative caspase 1 activity from primary patient leukemia samples *TP53* wild-type and mutant treated with (J )TNFα (50ng/mL) (n=5 *TP53^wt^*, n=6 *TP53^Mut^*) and (K) doxorubicin (1µM) (n=3 each group, in triplicate). *p≤.05.

NLRP1 is involved in forming the inflammasome complex that upon activation can lead to caspase 1 activation by protease activity and pyroptosis, an inflammatory form of cell death (29). In a mutagenesis screen for neutropenia, a hyperactive *Nlrp1a* allele was identified in mice to result in an inflammatory state and HSPC depletion (30). Thus, persistent activation of this pathway leads to cytopenias and wild-type HSPC loss. In the TCGA AML cohort as a univariate variable, elevated expression of *CASP1,* an interferon-responsive gene, is associated with worse prognosis, suggesting inflammatory states may be important in disease outcomes (**SFig. 5D**) (31,32). Adapting to this inflammatory stress may also determine leukemia HSPC evolution. As p53 is a regulator of many cell death pathways and the observed associated gene expression changes in our leukemia model, we tested whether p53 also regulates *NLRP1* expression. When we treated cell lines with MDM2 inhibitors (AMG232 and MS3227 250nM), activators of p53 signaling, we observed increased mRNA expression of *NLRP1* that also correlated with increased protein expression (**Fig. 7B-C** and **SFig. 5E**). We also treated leukemia cell lines with chemotherapy compounds, doxorubicin, 5FU, and cytarabine. As DNA-damaging agents, these treatments increased p53 and the p53 canonical target p21 *Cdkn1a* and *NLRP1* mRNA (**Fig. 7D** and **SFig. 5F**). This appears specific to *NLRP1* as *NLRP3* expression did not appreciably change (**SFig. 5G**). Thus, *NLRP1* is a target of TP53 activation in response to cellular stress.

### Loss of NLRP1 leads to relative resistance to pyroptotic stress

NLRP1 is a cellular stress sensor that can be activated through a variety of signals including pathogens, p38 kinase signaling and translational stalling (33–35). We demonstrated in vitro that treatment of WT mouse bone marrow cells with INFγ and TNFα can implicate this pathway with phosphorylation of p38 and caspase 1 activation resulting in cleavage of gasdermin D and also p53 signaling (**Fig. 7E** and **SFig. 5H**). Cleaved gasdermin D is required for multi-unit pore complex formation associated with pyroptosis (36). These cytokines also activated signaling to phosphorylate ZAKα, a sensor of stalled ribosomes and translation (**SFig. 5I**) (37). Likewise, treatment of leukemia cell lines with doxorubicin, an anthracycline class used in many regimens to treat acute leukemias, causes translational stress through ribosome stalling, and activates this pathway (**Fig. 7F**). To test the significance of NLRP1 activation, we generated knockout cells for *NLRP1* in leukemia cell lines and upon exposure to doxorubicin, cleavage of gasdermin D is reduced (**Fig. 7F**). Similarly, caspase 1 activity is diminished with doxorubicin treatment (**Fig. 7G**). Pyroptosis is a non-apoptotic cell death pathway and may in part contribute to anthracycline-induced cell death. *NLRP1* knockout cells that are deficient in pyroptosis demonstrate decreased sensitivity to doxorubicin (OCIAML3 IC50, CTRL 236nM, KO 332nM) (**Fig. 7H** and **SFig. 5J-L**).

To assess the relevance of this pathway in primary samples, we examined bone marrow cells derived from aged mice with myeloproliferative features (marked by splenomegaly), *Tet2^-/-^ Tp53^-/-^* and *Tp53^-/-^* mutant cells demonstrated decreased activation of pyroptosis with lower cleaved gasdermin D levels compared to *Tet2^-/-^*and a wild-type control (**Fig. 7I**). This was also seen in culture of bone marrow cells with inflammatory cytokines (**SFig. 5M**). To assess relevance of T53 signaling and pyroptosis in human leukemia samples, we examined primary AML samples that were either WT or mutant for *TP53.* In primary culture, we exposed patient mononuclear cells (MNCs) to the inflammatory cytokine TNFα for 24hr. *TP53^WT^* AML MNCs in general mounted a greater response of the inflammatory pathway with activated caspase 1 activity compared to *TP53^Mut^* MNCs (**Fig. 7J**). Similarly, treatment with doxorubicin also elicited higher caspase 1 activity in *TP53^WT^* AML MNCs compared to *TP53^Mut^* (**Fig. 7K**). Thus, the p53-NLRP1 pathway may be important for stress and chemotherapy response, and its loss would provide a growth advantage to leukemia cells by decreasing the likelihood of pyroptotic cell death.

## Discussion

*TET2* mutations can drive CH and although they are commonly found in myeloid malignancies such as AML and MDS they are also present in lymphoid leukemias and lymphomas (38). In *TET2* mutant CH, inflammation is common and associated with aging associated disease. In animal models of *TET2* LOF mutations, activation of the inflammasome has been previously observed (39). Independently, cytokines, and particularly INFγ have been shown to inhibit stem cell renewal in WT hematopoiesis (40). Similarly, a TNFα response pattern has been shown to confer advantage to *Tet2* mutant cells compared to wild-type cells *in vivo* (41). A normal function of inflammation and cytokines such as TNFα is to promote myelopoiesis during acute infections to generate mature monocytes and granulocytes but results in loss of select HSPCs (42). In the chronic condition this may promote myeloproliferative states and continued disruption of normal HSPC function in which additional mutations can cooperate to confer even further advantage. Thus, *TET2* mutation enhances HSPC fitness in an inflammatory stressed environment that shapes bone marrow dynamics. Despite *TET2-*mutant HSPC advantage and tolerance in this inflammatory context, this environment still represents a stress that may select for additional genomic events leading to transformation.

In advanced hematologic malignancies, additional mutations must co-occur with *TET2* mutations to lead to overt disease. *TP53* is one such mutation, regulating key cell death pathways. This is also true of *TP53* mutant CH acting as the founding clone of more advanced leukemias. *TP53* mutations can be selected in the setting of genotoxic stress such as with chemotherapy (43). However, secondary effects of chemotherapy on tumor microenvironment inflammation may also contribute the selection of *TP53* mutations (44).

Our model of *Tet2* and *Tp53* mutation demonstrates both lymphoid and myeloid disease potential. The combined mutations generate a competitive stem cell growth advantage compared to wild-type and single *Tet2* or *Tp53* mutant stem cells and evolution to disease can occur with either founding mutation. Chromosomal instability (CIN) and subsequent transformation develop when these mutations are both present. CIN is characteristic of *TP53* mutant hematopoietic malignancies and may dictate disease outcome. These processes may be linked as CIN is known to be associated with activated cell-intrinsic inflammation (45).

In this manuscript, we characterized the *Tet2-Tp53* double mutant acute leukemia phenotype and identified elevated IFNγ and TNFα pro-inflammatory cytokine production and signaling during lymphoid leukemic transformation. This environment creates an advantage for malignant cells. The *Tet2-Tp53* double mutant stem cells have increased self-renewal activity compared to wild-type and single mutant cells when exposed to IFNγ in particular. This cell intrinsic advantage exists at pre-malignant stage, allowing for increased inflammation and outgrowth of the mutant clone.

We further identified a *TP53*-dependet *NLRP1–*mediated inflammasome pathway activation as a contributor to disease progression. An investigation of TP53 target genes by meta-analysis of over 300 studies proposed *NLRP1* as a high-confidence target amongst 343 genes (46). TP53 has also been shown to activate the inflammatory cell death pathway in solid tumors (47). TP53 directly regulates *NLRP1* expression, and an NLRP1-mediated pathway is utilized in stress sensing. In particular, the expression of inflammatory cytokines can activate ZAKα, the signaling activator of this response (37). This pathway is also important for understanding treatment response to chemotherapy, secondary to ribosome stalling from disrupted translation. Cytokine stimulation, especially with IFNγ and TNFα , poses a stress to hematopoietic stem cells and combined *Tet2* and *Tp53* provides greater HSPC tolerance. Loss of *TP53* renders cells more resistant to stresses that signal through *NLRP1,* including those involved in ribosome stalling and inflammatory stress. Understanding how hematopoietic stem cells evolve from a CH state to a leukemic state is important to identify ways to prevent or treat transformation. Inflammatory and IFNγ signaling gene signatures have also been associated with treatment resistance and worse prognosis in leukemia (48–50). We further suggest that targeting signal transduction pathways from these cytokines may represent a therapeutic strategy as these factors may sustain leukemia (51). Alternatively, in mutant leukemias that demonstrate inflammatory signatures and retain inflammatory cell death pathway components, the pyroptosis pathway is of potential interest in therapy as small molecule activators of NLRP1 have shown anti-leukemic activity (52,53). Taken together, our data suggest the clinical importance of these pathways in high risk leukemias with *TET2 TP53* co-mutations, and underscore the potential for targeting inflammation which arise from pre-existing CH.

## Materials and Methods

### Primary genetically engineered mouse models husbandry and treatments

The conditional C57BL6/J *Mx1-Cre^+^* (Jax strain # 003556) *Tet2*^fl/fl^ (Jax strain # 017573) and germline *Tp53^-/-^*(Jax strain #002101) mice have been previously described individually (18,54). They were crossed together to generate *Mx1-Cre^+^ Tet2*^fl/fl^ *Tp53^-/-^* mice. Cre expression was induced by intraperitoneal injections (IP) of poly(I:C) (Invivogen) every other day at a dose of 20 μg/g for 3 doses. Rosa 26*^TdTomato^* (Jax strain #007909), *Jak1^fl/fl^, Tp53^fl/fl^ and Scl-CreER^T^* (Jax strain # 037467) mice were crossed into the *Tet2*^fl/fl^ strain as indicated and treated with Tamoxifen to induce Cre recombination. Routine genotyping primers are in STable 4.

All animal procedures were conducted in accordance with the Guidelines for the Care and Use of Laboratory Animals and were approved by the Institutional Animal Care and Use Committees at Memorial Sloan Kettering Cancer Center (MSKCC) and the Icahn School of Medicine at Mount Sinai.

### Histology and hematology assays

Murine tissue samples were fixed in 4% paraformaldehyde and embedded in paraffin. Sections were mounted on slides before hematoxylin and eosin staining. Reticulin staining was done with an ammoniacal silver protocol. Peripheral blood smears were air dried, fixed in methanol, and stained with Wright-Giemsa Stain. Complete Blood Counts (CBCs) were acquired on an IDEXX ProCyte machine.

### Bone marrow transplantation and cell harvesting, drug treatment

Dissected femurs and tibias were isolated. BM was flushed with a syringe into RPMI 10% FCS media or centrifuged at 8,000 rpm for 1 minute. Spleens were isolated and single-cell suspensions were made by mechanical disruption using glass slides. Cells were passed through a 70-μm strainer. Red blood cells were lysed in ammonium chloride–potassium bicarbonate (ACK) lysis buffer. A total of 1 × 10^6^ cells were transplanted via tail-vein injection into lethally irradiated (950 Rad) C57BL/6 (Jax strain # 000664) or CD45.1 (Jax strain # 002014) host mice. For Ruxolitinib (Chemietek) treatment, mice were treated at 60mg/kg daily (in 20% captisol) by oral gavage. Mice were randomized by WBC count prior to treatment. Treatment was unblinded. For LPS (Sigma) treatment, mice were treated at 2.5mg/kg twice weekly for 3 weeks. (* p<.05, ns-not significant).

### RNA Sequencing and Analysis

Cell populations were sorted using a Sony SH800 and RNA stored and isolated using TRIzol LS and RNeasy/AllPrep (Qiagen) respectively. After RiboGreen quantification and quality control by Agilent BioAnalyzer, 0.8-1 ng total RNA with RNA integrity numbers ranging from 7.5 to 9.9 underwent amplification using the SMART-Seq v4 Ultra Low Input RNA Kit (Clonetech catalog # 63488), with 12 cycles of amplification. Subsequently, 6 ng of amplified cDNA was used to prepare libraries with the KAPA Hyper Prep Kit (Roche 07962363001) using 8 cycles of PCR. Samples were barcoded and run on a HiSeq 4000 in a PE50 run, using the HiSeq 3000/4000 SBS Kit (Illumina). An average of 26 million paired reads were generated per sample and the percent of mRNA bases per sample ranged from 57% to 70%. Aligned RNA was analyzed for fold change. FASTQ files were mapped, demultiplex, and transcript counts were enumerated using STAR. Counts were input into R and RNA-sequencing analysis was completed using DESeq2. Differentially expressed genes were identified with an adjusted *p* value of 0.05. Gene set enrichment analysis was performed using the GSEA package with gene sets extracted from the Msigdbr package.

### ERRBS and Differential Methylation Analysis

Cell populations were sorted using a Sony SH800 and genomic DNA was digested with MspI. DNA fragments were end-repaired, adenylated, and ligated with Illumina kits. Library fragments of 150–250 bp and 250–400 bp were isolated. Bisulfite treatment was performed using the EZ DNA Methylation Kit (Zymo Research). Libraries were amplified and sequenced on an Illumina HiSeq. Differential methylation analysis was performed on the resulting ERRBS data using methylKit for finding single nucleotide differences (*q*-value < 0.01 and methylation percentage difference of at least 25%) and eDMR for finding differentially methylated regions (55). Gene annotations were carried out by overlapping the CpG sites and regions with the exons, introns, UTR5, UTR3 and 5-kb upstream regions of RefSeq genes for genome assembly mm10.

### Mouse exome sequencing

DNA was isolated from bone marrow tissues using Qiagen DNA extraction kit. After PicoGreen quantification, 100 ng of mouse genomic DNA were used for library construction using the KAPA Hyper Prep Kit (Kapa Biosystems KK8504) with 8 cycles of PCR. After sample barcoding, 90-220 ng of each library were pooled and captured by hybridization with the mm_IMPACT (Nimblegen) (Integrated Mutation Profiling of Actionable Cancer Targets) assay, which captures all protein-coding exons and select introns of 578 cancer-related genes. Capture pools were sequenced on the HiSeq 4000 or NovaSeq 6000, using the HiSeq 3000/4000 SBS Kit or NovaSeq 6000 Reagent Kit (Illumina), respectively, for PE100 reads. Following these criteria, the average coverage was 361X for tumor samples and 191X for normal controls.

Library sequences then underwent standard analysis pipeline to remove adapter sequences and aligned to the mouse reference genome using BWA. Sequences were processed to identify unique reads then analyzed by a somatic mutation calling program (MuTect2). Copy number determination was compared to a pooled normal library using genome segment binning with an analysis pipeline previously described using Varbin algorithm (56). The analysis calculates a log ratio between sample and reference.

### CRISPR targeting

Guide oligonucleotides were purchased from IDT. Complementary oligonucleotides were annealed and ligated LentiCRISPRv2 puro (Addgene 98290). Editing was confirmed by PCR and sequencing of genomic DNA including the target site. Sequences used for targeting and sequencing are in STable 4.

### Lentivirus production

For lentivirus production, plasmids were co-transfected into 293T LX cells with the packaging vector psPAX2 (Addgene 12260) and the envelope vector pMD2.G (Addgene 12259) using the jetPRIME transfection reagent (Polyplus) following the manufacturer’s protocol. Media was changed at 24-hours post transfection. Viral-containing supernatant was collected 72 hours after transfection and passed through a 0.45-micron filter. The virus was ultracentrifuged at 24,000 rpm for 2 hours at 4°C using a Sorvall Ultracentrifuge. Viral pellets were resuspended in serum free medium, aliquoted and stored at −80°C for long-term storage.

### In vitro Colony-Forming Assays

Bone marrow cells (30,000 per assay well) were set into cytokine-supplemented methylcellulose medium (Methocult, M3434; STEMCELL Technologies). Colonies propagated in culture were scored at day 7-10. Cells were collected again with PBS washout and re-plated using similar conditions for stem-cell in vitro re-plating potential. Recombinant mouse IFNγ and TNFα (BioLegend) used at 50ng/mL.

### Flow Cytometry and FACS

Antibody staining and flow cytometry analysis was performed as previously described (18). Cells were lysed in ACK lysis buffer except for CD71 Ter119 staining. Stem cell enrichment was performed using the Progenitor Cell Enrichment Kit (Stem Cell Technologies #19856). For peripheral blood and bone marrow analysis, BioLegend antibodies clones were for CD3 (17A2), CD45R/B220 (RA3-6B2), Gr-1 (RB6-8C5), CD11b 450 (M1/70), CD45.2 (104), CD117 (2B8), CD45.1 (A20), Sca-1 (D7), FcγRII/III (2.4G2), CD34 (RAM34), CD71 (RI7217), Ter119 (TER-119), CD150 (9D1), and CD48 (HM48-1). Live and dead cell discrimination was performed using DAPI and PI stains.

### Quantitative Real-Time PCR

RNA was extracted using Qiagen RNAEasy kits. cDNA was then reverse transcribed using ProtoScript II First Strand cDNA synthesis kit (New England Biolabs) per the manufacturer’s protocol. qPCR was performed using a SyberGreen marker and Applied Biosystems Sybr Master Mix. Readings were acquired on an Applied Biosystems Real-Time PCR instrument. The quantity of cDNA was normalized according to the expression of *Actin* and *GAPDH* Data. The delta–delta Ct ratio technique was used to normalize for the expression of these housekeeping genes. For human AML samples, genomic DNA was extracted from frozen mononuclear samples using Qiagen DNA extraction kit. Relative DNA copy was determined through qPCR of *NLRP1* and *TP53* relative to an internal control gene (*IFNB1*). Primers sequences are in STable 4.

### Cell Culture and treatments

THP-1 (CVCL_0006), MV4-11 (CVCL_0064), OCI-AML3 (CVCL_1844), MOLM13 (CVCL_2119) cells were cultured in RPMI 1640 medium. HEK293T (CVCL_0063) and HCT 116 (CVCL_0291) cells were cultured in DMEM. All cell lines were cultured in a medium containing 10% FBS and 1% penicillin/streptomycin at 37°C with 5% CO_2_. Primary murine cells for short term cytokine induction were cultured in IMDM 2% FBS. Primary AML cells were then seeded at 5×10^4^ cells/mL density in STEMSPAN II media mixed with the following reagents: recombinant human SCF, FLT3L, and TPO at 20 ng/mL. Doxorubicin, AMG 232 (250nM) was supplied by MedChemExpress. Puromycin was used at 1–2μg/mL. MS3227 PROTAC (250nM) was used as previously described (57). Recombinant INFγ and TNFα (Biolegend) used as indicated. Cell viability and proliferation was performed using the Promega CellTiter-Glo Luminescent Cell Viability Assay.

### Cytokine Quantification

Cytokine quantification was performed with Eve Technologies (Calgary, AB) using the Mouse Cytokine 44-Plex Discovery Assay using Luminex and multiplexed laser bead capture.

### Caspase 1 Assay

Cells were collected from cultures and assayed for caspase 1 activity using the Promega Caspase-Glo 1 Assay according to manufacturer’s instructions and data collected after signal stabilization. Luminescence was determined using a SpectraMax Spectrofluorometer. Vehicle treated cells were set to 100 and treated cells’ activity relative to this level.

### Western blot analysis

Protein lysates were prepared with cell lysis buffer with protease and phosphatase inhibitors, and protein concentrations were determined using Bradford assays. Proteins were run on NuPAGE Bis-Tris gels and transferred to nitrocellulose or PVDF membranes. Membranes were blocked in 5% non-fat milk in 1X TBST and incubated with respective primary antibodies overnight at 4°C (1:1000 dilution), followed by an anti-rabbit (CST), or anti-mouse (CST) secondary IgG HRP antibody. Protein bands were then detected using chemiluminescence-detection and the signal was visualized using a digital imager (BioRad ChemiDoc). NLRP1 (CST, E3F2U, #56719), p21 (human) (CST, 12D1, #2947) , GAPDH (CST, D16H11, #5174), Cleaved Gasdermin D (human) (CST, E7H9G, #36425), Cleaved Gasdermin D (mouse), Gasdermin D (CST, E9S1X, #39754), Vinculin (CST, E1E94, #13901), phospho p38 (CST, D3F9, #4511), p53 (CST, D2H9O, #32532), p21 (mouse) (CST, F2C7C, #39256), phospho (S165)-ZAK (Bioworlde, polyclonal, #BS64561).

### Statistical Analyses

Statistical analyses were performed using the one and two tailed Student *t* test using GraphPad Prism (GraphPad Software, San Diego, CA) unless otherwise noted. *p* ≤0.05 was considered statistically significant. Drug response tests were performed using non-linear fit and sum-of-squares F-test in GraphPad Prism.

### Patient AML Data

We used patient datasets of acute leukemia patients available from the cBioPortal for cancer genomics analysis of co-occurring mutations, survival, and variant allele frequencies.

### Patient Samples

Patients at the Mount Sinai Hospital provided informed consent for biological samples research, in accordance with the Declaration of Helsinki. Primary AML cells were obtained through the Hematological Malignancies Tissue Bank (HMTB) of Tisch Cancer Center through an IRB-approved protocol. Both male and female adult patients greater than 18 years of age were included as potential subjects.

### Data availability

Sequencing data are deposited at Gene Expression Omnibus GSE300000, GSE305288, GSE305295.

## Acknowledgements

Alan Shih is supported by grants from the Leukemia Research Foundation, the American Society of Hematology (ASH) Scholar Program, and the Gilead Sciences Research Scholars Program.

Research was supported by the Tisch Cancer Center - Cancer Center Support Grant from the NCI (P30 CA196521). We acknowledge the use of the Integrated Genomics Operation Core, funded by the NCI Cancer Center Support Grant (CCSG, P30 CA08748), Cycle for Survival, and the Marie-Josée and Henry R. Kravis Center for Molecular Oncology.

We thank the Epigenomics Core at Weill Cornell Medicine for methylation studies and Nicholas Socci of the Bioinformatics Core at the Sloan Kettering Institute for exome sequencing analysis.

