## Supplemental Figures for "*TET2* and *TP53* Mutations Cooperatively Modulate the Response to Inflammation to Promote Leukemic Transformation"

#### Supplemental Fig. Legends

**SFig. 1. *Tet2 Tp53* GEMM development and extended phenotype**. **A** Genotyping for *Tp53* deletion (del) and $Tp53^{wt}$ (+) and *Mx1-cre* (+ vs -) from tail tissue and for deletion of *Tet2* in the peripheral blood following poly(I:C) (PIPC) administration and transplantation. floxed (fl), deleted (del). **B**, Flow cytometry gating for B220 dim fraction (immature B-cells). **C**, Peripheral blood fraction of myeloid ($CD11b^{+}$) vs B-lymphoid ($B220^{+}$) cells in recipients from 2 separate *Tet2 Tp53* mutant donors. **D**, Flow cytometry of peripheral blood from bone marrow transplanted recipient mice (n=4) with lymphoid (top) or myeloid (bottom) *Tet2 Tp53* mutant disease cells, compared to a wild-type control.

**SFig. 2. *Tet2 Tp53* disease leads to chromosomal instability (CIN)**. Copy number analysis with exome sequencing compared to normal in individual diseased mice. **A**. *Tp53* single mutant disease without significant copy number changes. **B.** Myeloid disease with evidence of copy number changes. **C**. *Tet2 Tp53* lymphoid leukemia from a separate transplant recipient as in (Fig. 5). **D-F.** *Tet2 Tp53* Lymphoid disease from separate primary mice.

**SFig. 3. Differential gene-expression in *Tet2 Tp53* double mutant leukemic cells versus controls**. **A**, Volcano plot of gene expression *Tet2 Tp53* double mutant leukemic cells versus controls (expression fold change vs adjusted p-value). Select individual genes highlighted – upregulated in orange, downregulated in purple. **B**, GSEA expression analysis and plot of significant gene signatures q<0.05, NES >1.2. (Arrow) Hallmark interferon gamma signature. **C**, qPCR relative gene expression of *Ifng* and target genes *Casp1* and *Stat1* in WT compared to *Tet2 Tp53* double-mutant leukemic bone marrow. **D**, Reticulin staining in wild type, *Tet2 Tp53* mutant disease, and non-inflammatory *Tet2 Flt3* double-mutant leukemic bone marrow.

**SFig. 4. Cytokine response in colony formation and induction in *Tet2 Tp53* mutant cells**. **A&B**, Colony forming assay with wild type, *Tet2, Tp53*, and *Tet2 Tp53* double-mutant bone marrow cells. Re-plating assays treatment with (A) vehicle and (B) IFNγ and TNFα (50ng/mL). For the first plating with cytokine, percentage of immature -granulocyte, erythrocyte, monocyte, and megakaryocyte CFU-GEMM quantified. **C**, qPCR of the relative gene expression of *Ifng* and *Tnfa* following *in vitro* culture induction of whole bone marrow cells with LPS (100ng/mL).

**SFig. 5. Inflammation response associated with *TP53* mutation**. **A**, qPCR of *Nlrp1a* relative gene expression induced by IFNγ (50ng/mL) treatment of whole bone marrow cells in culture at 4hrs. **B**, qPCR of *Nlrp1a* relative gene expression in wild type, *Tet2, Tp53*, and *Tet2 Tp53* double-mutant bone marrow cells. **C,** qPCR of relative copy number (normalized to internal control gene) of *TP53* and *NLRP1* genomic DNA derived from myeloid malignancy MNC samples with *TP53* wild-type and *TP53* del 17p samples. **D**, Overall survival of the TCGA AML cohort divided into *CASP1* highest (red) n=53 and lowest (blue) n=53 expressing AMLs. **E**, Western blot expression of NLRP1, p21 and Vinculin in HCT116 cells treated with MDM2 inhibitor MS3227 1μM. **F&G**, qPCR relative expression of (F) *CDKN2A* and (G) *NLRP3* in MV4-11 cells treated with vehicle or doxorubicin (1μM), 5FU (25μM), or cytarabine (25μM). **H&I**, Western blot expression of (H) phospho-p38, GAPDH, cleaved Gasdermin D, p53, p21, and (I) phospho-ZAK, vinculin in mouse wild-type bone marrow (BM) samples and THP-1 cell line treated with IFNγ, TNFα, IFNγ + TNFα (50ng/mL), or Anisomycin (1μM). **J&K**, Dose response curves to doxorubicin in (I) MV4-11 and (J) THP-1 cells, control and *NLRP1* knockout. Curves analyzed by variable slope least squares fit. Below, Sanger sequencing demonstrating editing at *NLRP1* locus (arrow). Gray bar indicating quality of read that declines subsequent to CRISPR editing site. **L,** Western blot of cleaved Gasdermin D and vinculin in MV4-11 control and *NLRP1* knock out cells treated with Doxorubicin (1μM). **M**, Western blot of cleaved Gasdermin D and vinculin in wild type, *Tet2, Tp53*, and *Tet2 Tp53* double-mutant cells bone marrow cells treated with vehicle or IFNγ (50ng/mL at 4hr and 24hr). *p<.05, ns- not significant.

Supplemental Figure 1

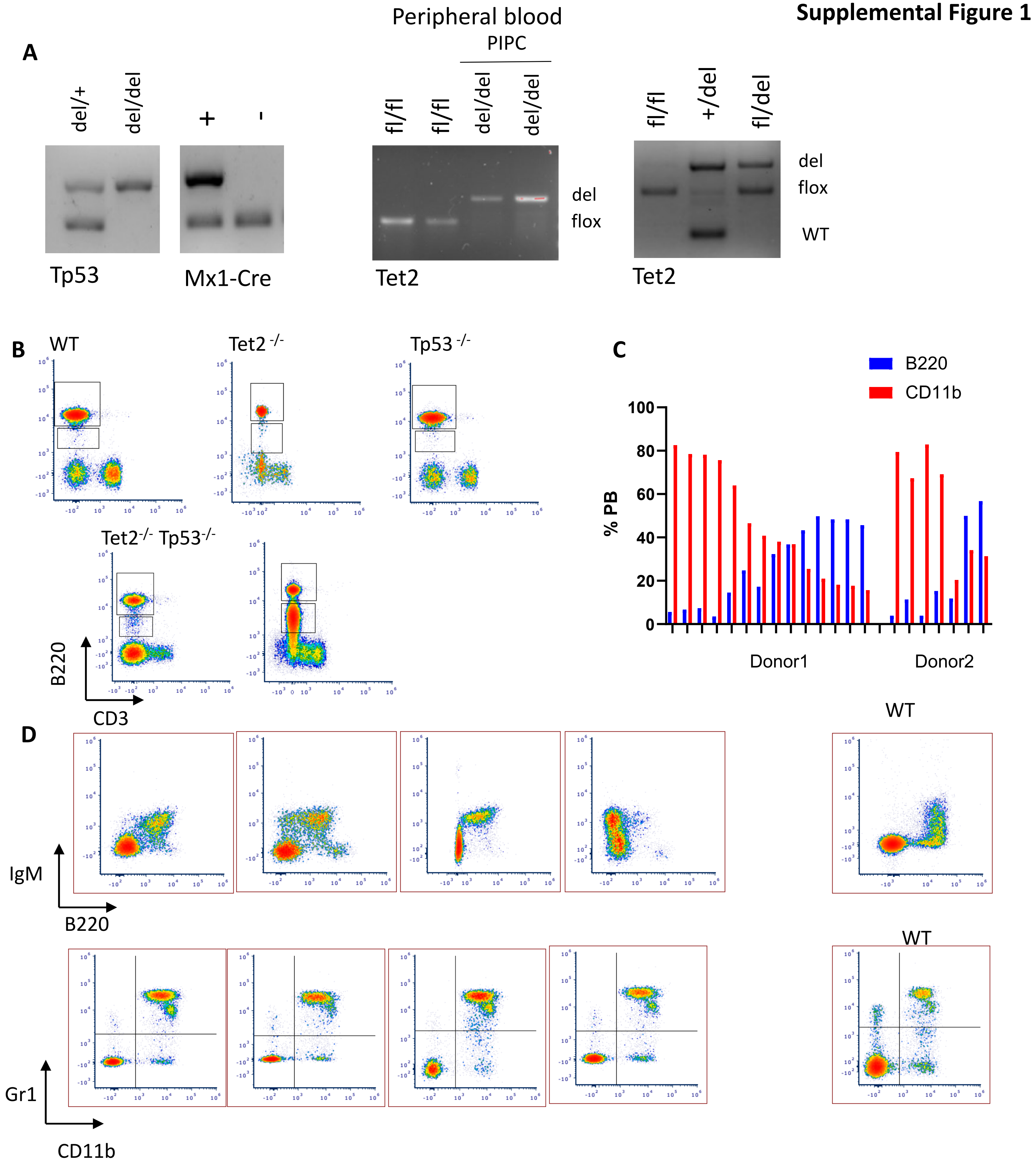

**Supplemental Figure 2**

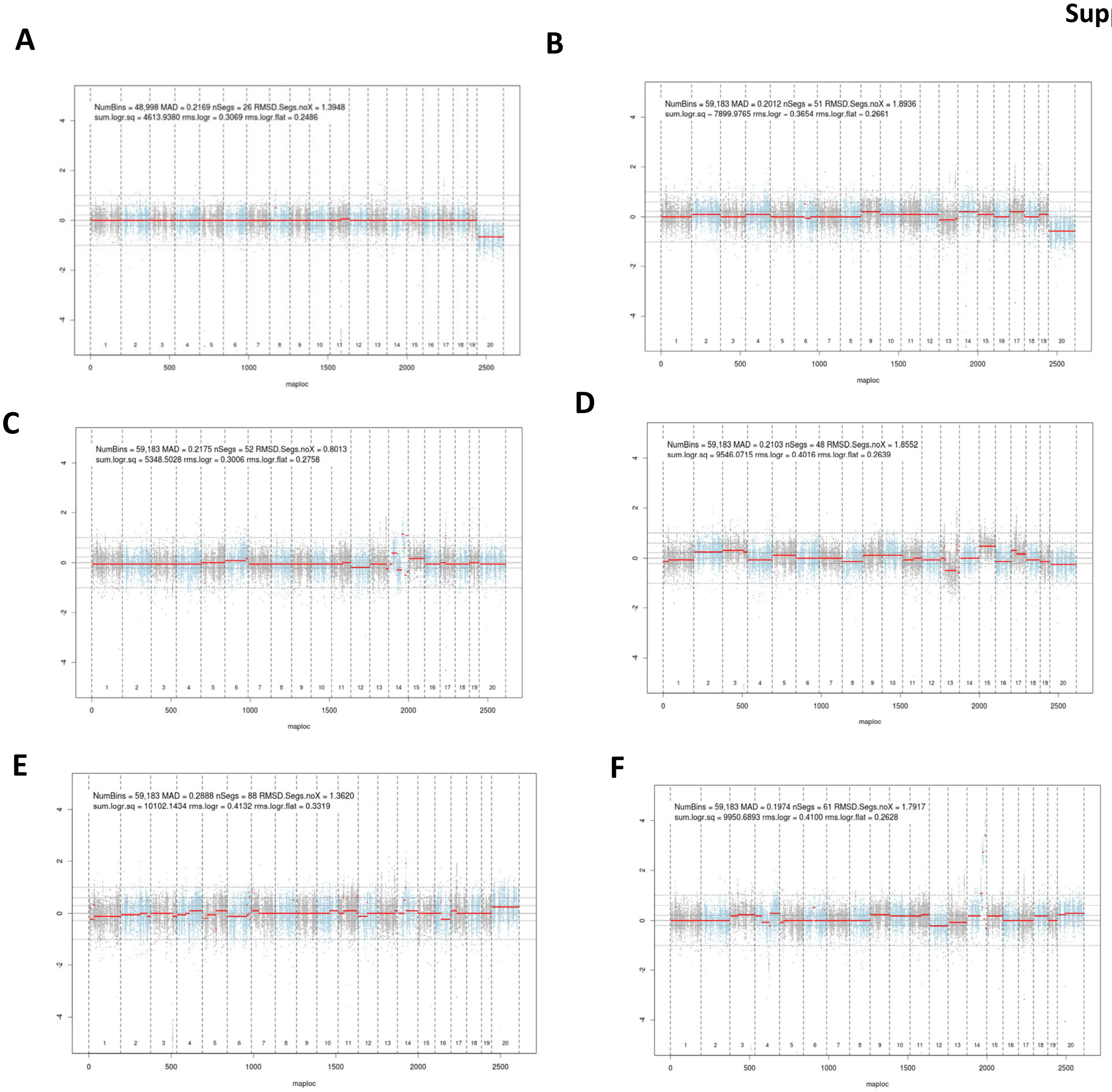

#### Supplemental Figure 3

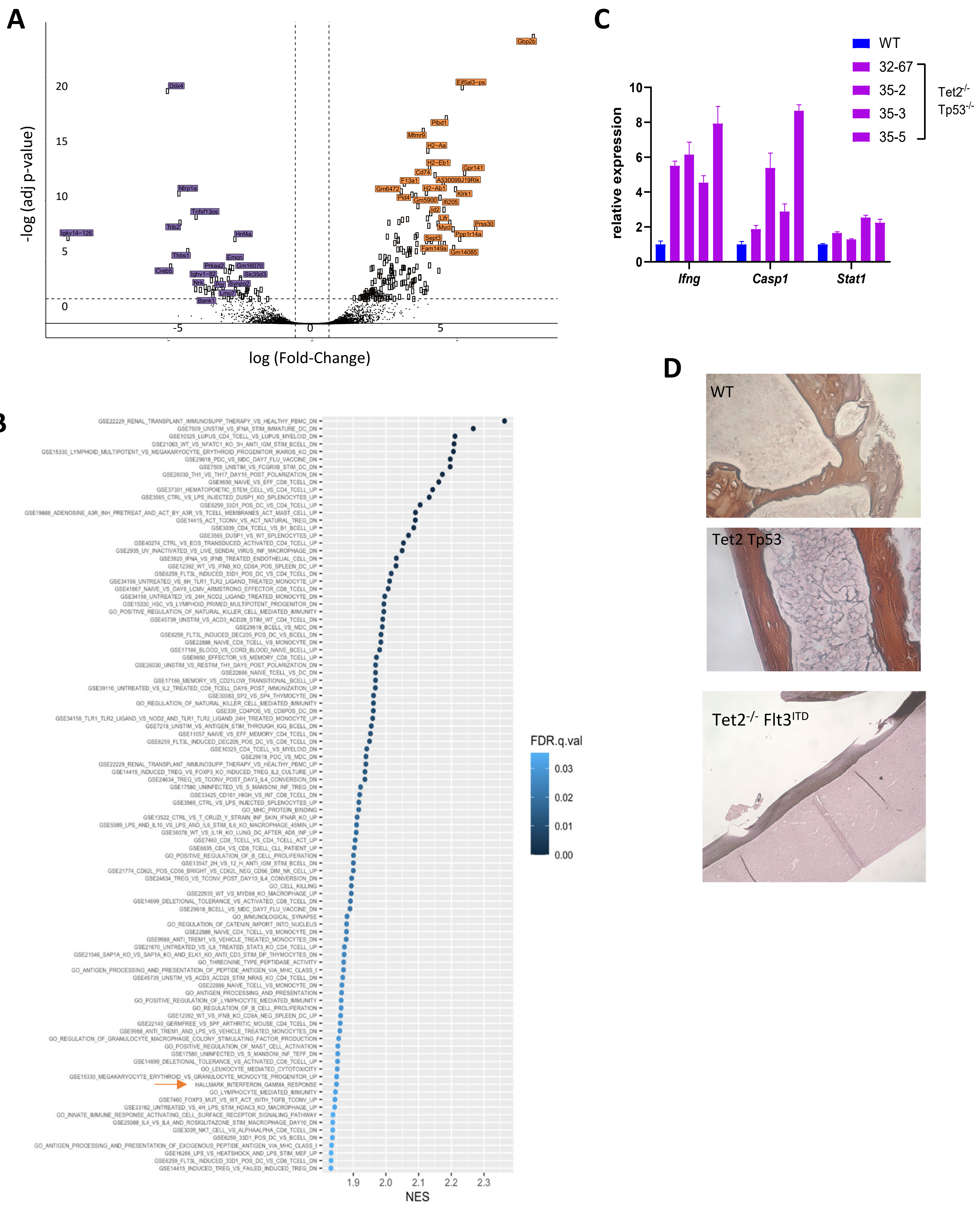

### Supplemental Figure 4

**A**

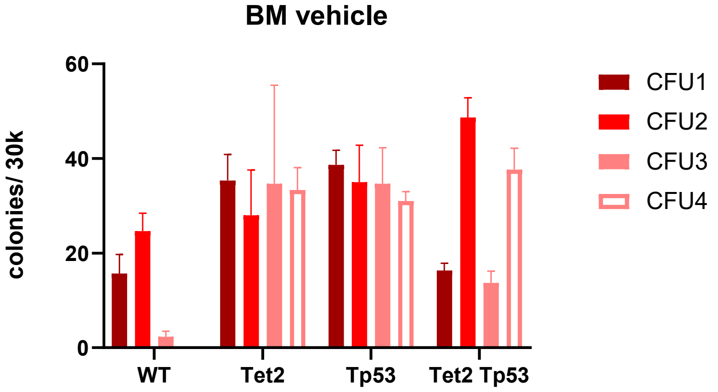


**B**

IFNγ and TNFα (50ng/mL) CFU assay

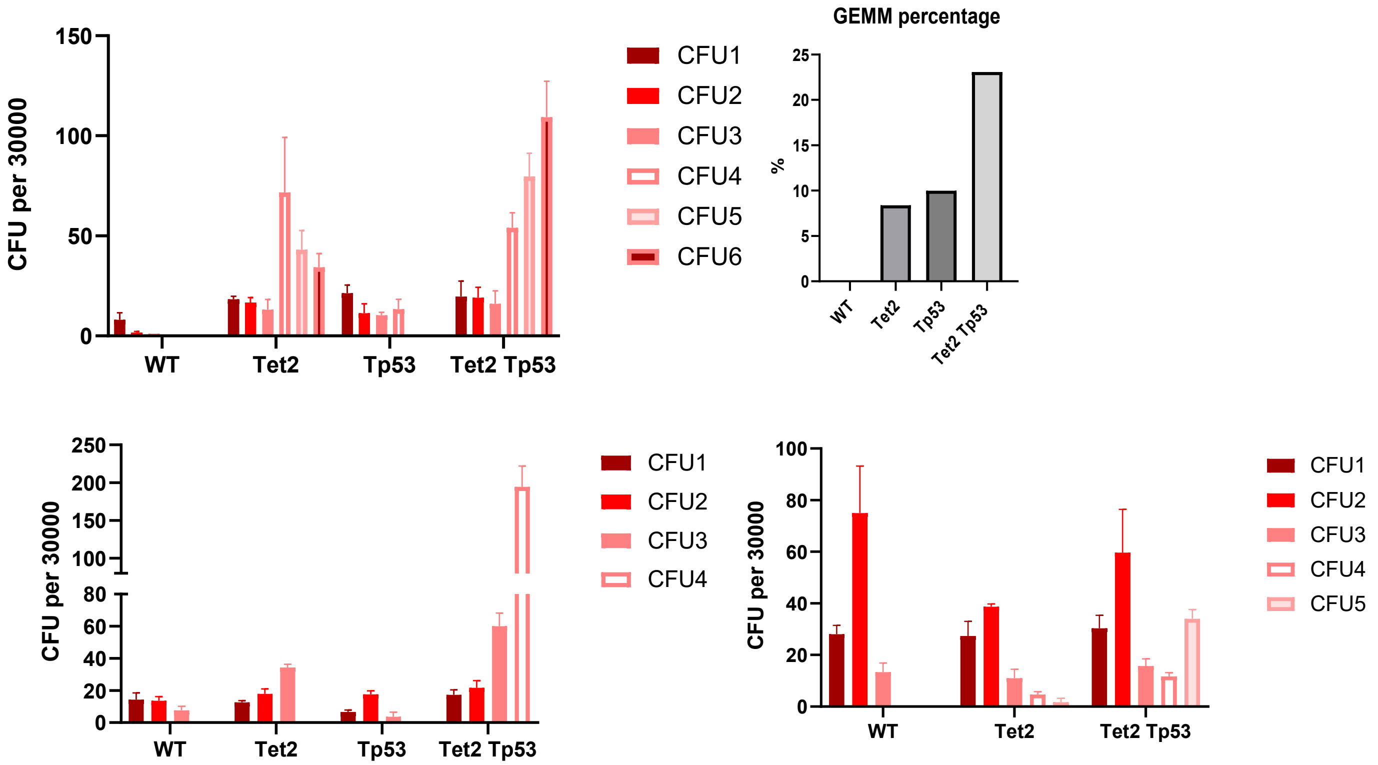


**C**

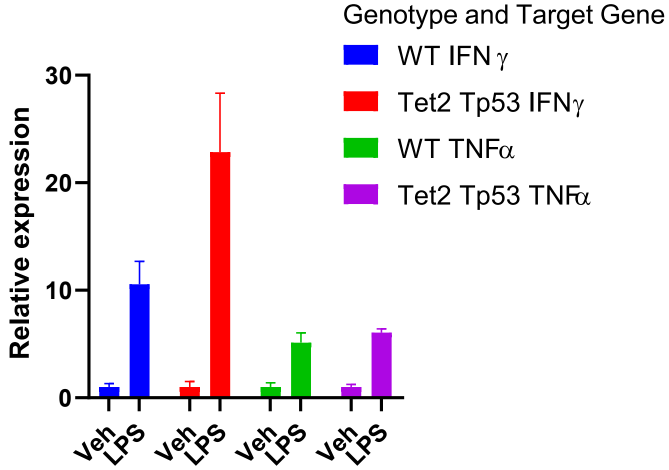

**Supplemental Figure 5**

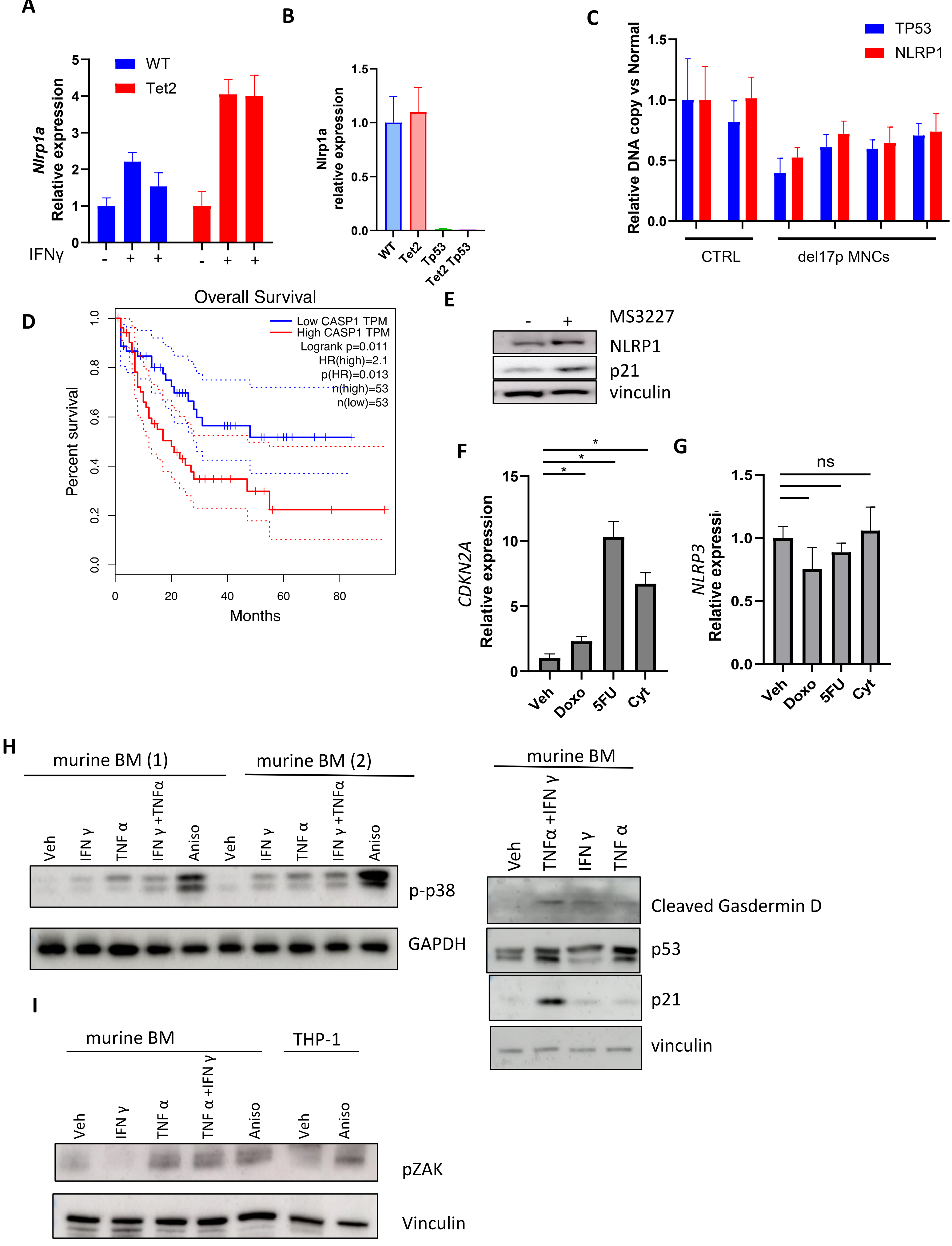

### Supplemental Figure 5 con't

J

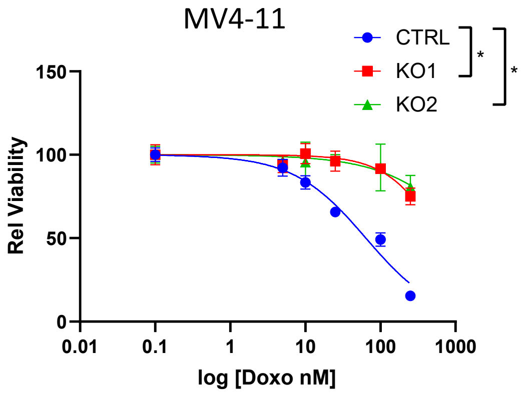


K

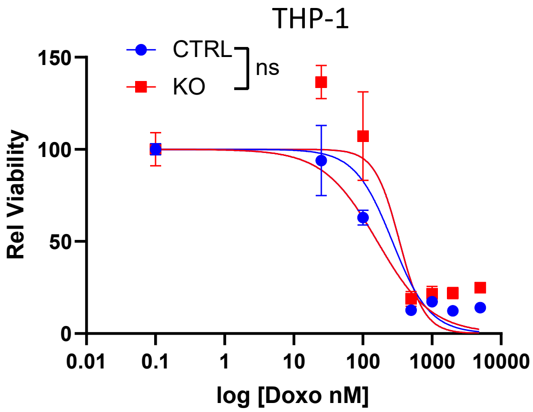


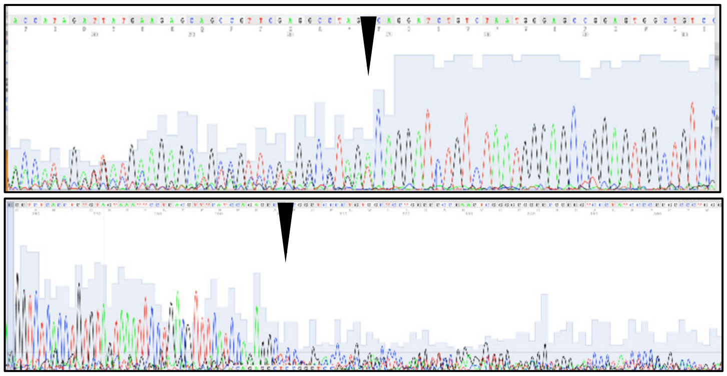

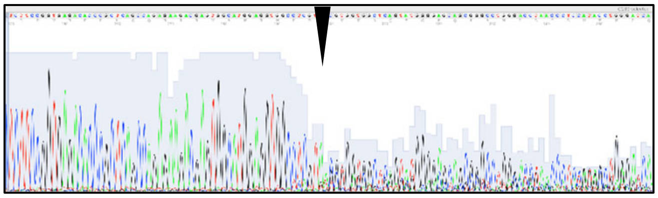

L

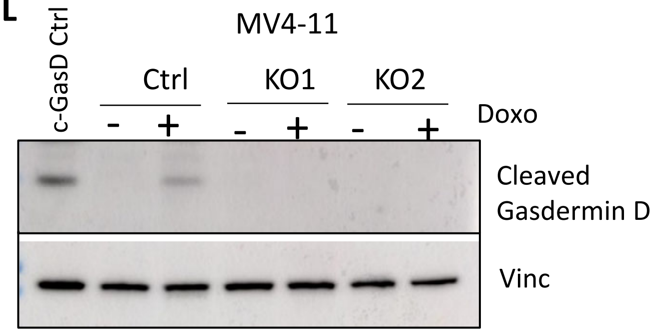


M

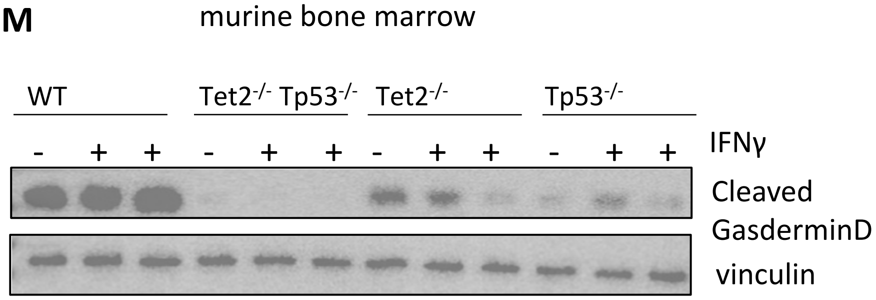
